# Estimating the replicability of Brazilian biomedical science

**DOI:** 10.1101/2025.04.02.645026

**Authors:** The Brazilian Reproducibility Initiative, Olavo Bohrer Amaral, Clarissa França Dias Carneiro, Kleber Neves, Ana Paula Wasilewska Sampaio, Bruna Valério Gomes, Mariana Boechat de Abreu, Pedro Batista Tan, Gabriel Paz Souza Mota, Ricardo Netto Goulart, Nathalia Raquel de Souza Fernandes, Jimmy Hayden Linhares, Adriana Mércia Guaratini Ibelli, Adriano Defini Andricopulo, Adriano Sebollela, Adryano Augustto Valladão de Carvalho, Airton Pereira e Silva, Alana Silva Oliveira Souza, Alessandra Catarina Chagas de Lima, Alessandra Mara de Sousa, Alexander Birbrair, Alexandre Urban Borbely, Aline Maria Machado, Alinne do Carmo Costa, Aliny Pereira de Vasconcelos, Alvaro Henrique Bernardo de Lima Silva, Amanda de Souza, Ana Beatriz Rezende Paula, Ana Carolina Caetano Nunes, Ana Maria de Lauro Castrucci, Ana Paula Almeida Bastos, Ana Paula Farias Waltrick, Ana Paula Herrmann, Andre Gustavo Ferreira de Macedo, Andrelson Wellington Rinaldi, André Souza Mecawi, Andrés Delgado Canedo, Anna Paula Marçal, Anna Paula Perin Vidigal, Antonio Martins Monteiro, Beatriz Gonçalves Silva Rocha, Bianca Cruz Pachane, Brenda da Silva Andrade, Bruna Carla Casali, Bruno Popik, Camila Pasquini de Souza, Camila da Silva Dos Santos, Camilla Mendes Gonçalves, Camilla Ribeiro Barroso do Nascimento, Carla Pires Veríssimo, Carlos Eduardo Neves Girardi, Carmen Penido, Carolina Batista, Carolina Panis, Carolina Saibro-Girardi, Carolina da Silva Gouveia Pedrosa, Caroline de Carvalho Picoli, Cristiane Regina Guerino Furini, Daniel Pens Gelain, Daniel Sturza Lucas Caetano, Daniela Costa Silva, Daniele Cristina de Aguiar, Demetrius Antonio Machado De Araújo, Deoclécio Alves Chianca Júnior, Dilza Balteiro Pereira de Campos, Débora Aguirre Gonçalves, Débora Santos da Silva, Eduarda Godfried Nachtigall, Eloiza Lopes Lira Tanabe, Erika Seki Kioshima, Fabio Rodrigues Ferreira Seiva, Fabrício de Araujo Moreira, Felipe Saceanu Leser, Felipe Vanz, Fernanda Cacilda dos Santos Silva, Fernanda Magalhães Ferrão, Fernanda Nogueira Lotz, Flavia Fonseca Bloise, Flavia Regina Souza Lima, Flávio Alves Lara, Franciana Aparecida Volpato, Francieli Moro Stefanello, Francisca Nathalia de Luna Vitorino, Francisco Noé Fonseca, Fábio Jorge Moreira da Silva, Fábio de Almeida Mendes, Gabriel Vasata Furtado, Gilda Angela Neves, Giovanna Zanetti, Giulia Scarcella Cancelliero, Graziele Fernanda Deriggi Pisani, Guilherme Curi Aiub Casagrande, Gustavo Roberto Villas-Boas, Heitor Roque Oliveira Alvez da Cruz, Helena Lobo Borges, Heloisa Sobreiro Selistre De Araujo, Helvécio Cardoso Corrêa Póvoa, Hilana dos Santos Sena Brunel, Hugo Bayer, Igor Luchini Baptista, Isabel Werle, Isabela Alcântara Barretto Araújo Jardim, Isabela Aparecida Divino, Janaina Menezes Zanoveli, Jane de Oliveira Peixoto, Jaqueline de Carvalho Rinaldi, Jeane Bachi Ferreira, Jeferson Luis Franco, Jeronimo Marteleto Nunes Rugani, Jociane de Carvalho Myskiw, Jorge Alberto Quillfeldt, José Marcos Janeiro Pereira da Costa, João Victor Roza Cruz, Julia Helena Oliveira de Barros, Julia Mello Barros, Julia Pinheiro Chagas da Cunha, Juliana Nunes Roson, Júlia Dummer Rodrigues de Freitas, Júlia Grigorini Mori Ayub, Karen Steponavicius Cruz Borbely, Karina Barbosa de Queiroz, Karina Dutra Asensi, Karoline dos Santos Rodrigues, Kellyne de Castro Vieira, Keyla Silva Nobre Pires, Kétlyn Talise Knak Guerra, Lays Adrianne Mendonça Trajano-Silva, Leandro Jose Bertoglio, Leonardo Vinicius Monteiro de Assis, Leticia Menezes Vasconcelos, Leticia Miranda Santos Lery, Letícia de Oliveira Marinho da Silva, Lorena de O Fernandes-Siqueira, Lucas De Oliveira Alvares, Lucas Gabriel Vieira, Lucas Henrique de Melo Falcione, Lucas Dos Santos da Silva, Lucianne Fragel Madeira, Luis Eduardo Duarte Nunes, Luiz Gustavo de Almeida Chuffa, Luiza Marques Prates Behrens, Marcos Vinicius Esteca, Maria Luiza Saraiva-Pereira, Maria Nathália Moraes, Maria Sueli Soares Felipe, Mariana Saldanha Viegas Duarte, Mariana Souza da Silveira, Marimelia A Porcionatto, Marisa Salvi, Maryana Albino Clavero, Mateus Lobato Ferreira, Matheus Alves de Moura, Matheus Gallas-Lopes, Mauro César Isoldi, Mayara Terra Villela Vieira Mundim, Michael Andrades, Micheline Freire Donato, Milena Piaia Barboza, Miliane Martins de Andrade Fagundes, Miriane Avelino Silva, Morgana Duarte da Silva, Munira Muhammad Abdel Baqui, Mychelle Pacheco de Souza, Mônica Corrêa Ledur, Nathalia Stark Pedra, Natália Iorio Lopes Pontes Póvoa, Natália Karla Bellini, Nauana Somensi, Nicolly Maria Payva Nunes, Nícia Pedreira Soares, Nívea Ferreira Silva, Pablinny Moreira Galdino de Carvalho, Paola Alejandra Cappelletti, Patricia Pestana Garcez, Patrícia Kellen Martins Oliveira Brito, Patrícia Rocha Martins, Paulo Roberto Moreira Lauar, Priscila Nicolicht-Amorim, Rafael Chitolina, Raiana Andrade Quintanilha Barbosa, Raul Alexander Gonzales Cordova, Regina Coeli Dos Santos Goldenberg, Rejane Giacomelli Tavares, Renata Krogh, Renata de Freitas Saito, Roberto Andreatini, Roberto Farina Almeida, Rodrigo Cunha Alvim de Menezes, Rodrigo Rorato, Roger Chammas, Rosangela Vieira de Andrade, Roselia Maria Spanevello, Rosiane Andrade Costa, Rubens Lima do Monte-Neto, Sabrina Alves dos Reis, Scheila Iria Kraus, Silvana Chedraoui Silva, Simone Michelan-Duarte, Stevens Kastrup Rehen, Sylvana Izaura Salyba Rendeiro de Noronha, Tania Maria Ortiga-Carvalho, Tanira da Silveira Prieto, Tatiane Renata Fagundes, Thabatta Karollynne Estevam Nakamura, Thadeu Estevam Moreira Maramaldo Costa, Tharcísio Citrângulo Tortelli Junior, Thays Barboza da Luz, Thiago Aparecido da Silva, Tiago Goss Dos Santos, Vanessa Beijamini, Victória Regina de Siqueira Monteiro, Vinícius Alexandre Wippel, Viviane Medeiros Oliveira Valença, Wanessa Fernanda Altei

## Abstract

Concerns over the replicability and reproducibility of published research have grown in many research fields, but empirical data to inform policies are still scarce. Biomedical research in Brazil expanded rapidly over the last three decades, with no systematic assessment of the replicability of its findings. With this in mind, we set up the Brazilian Reproducibility Initiative, a multicenter replication of published experiments from Brazilian science using three common experimental methods: the MTT assay, the reverse transcription polymerase chain reaction (RT-PCR) and the elevated plus maze (EPM). A total of 56 laboratories performed 143 replications of 56 experiments; of these, 90 replications of 45 experiments were considered valid by an independent committee. Replication rates for these experiments varied between 20 and 44% according to five predefined criteria. In median terms, ratios between group means were 58% lower in replications than in original experiments, while coefficients of variation were 82% higher. Effect size decrease was smaller for MTT experiments, original results with less variability and those considered more challenging to replicate, while t values for replications were positively correlated with researcher predictions about replicability, and negatively correlated with the rate of publications by the original article’s last author. Deviations from preregistered protocols were very common in replications, most frequently due to reasons inherent to the experimental model or related to infrastructure and logistics. Our results highlight factors that limit the replicability of results published by researchers in Brazil and suggest ways by which this scenario can be improved.

## Introduction

Concerns over the replicability and reproducibility of published results have increased in many areas of science over the past decade. Basic biomedical science has not been immune to these concerns: reports from the pharmaceutical industry (Begley & Ellis, 2012; Prinz et al., 2011) and prospective assessments of replicability in specific fields, such as cancer biology (Errington, Mathur, et al., 2021), spinal cord injury (Steward et al., 2012) and amyotrophic lateral sclerosis (Scott et al., 2008), have reported success rates ranging from 0 to 40%. Nevertheless, these projects have focused on selected subsets of papers in specific areas of science, and the dimension of the problem in the biomedical literature at large remains unknown.

Although concerns over reproducibility (defined here as reaching the same results when analyzing a set of data) and replicability (defined as finding similar results with new data) (National Academies of Science, 2019; Nosek et al., 2022), many of the proposed solutions to address them are local in scope, as institutions and funders are instrumental in setting up policies and incentives that can foster reproducible research (Munafò et al., 2017). Actionable data on the replicability and reproducibility of research produced within specific institutions, regions or countries, however, is extremely scarce.

Academic research in Brazil experienced rapid growth between the late 20^th^ and early 21^st^ century, driven by the expansion of higher education (Leta, 2012). Concurrently, the establishment of a national evaluation system for graduate education programs instituted new expectations for productivity, with a strong emphasis on journal-based metrics that has only recently begun to be reconsidered (Barata, 2019). More recently, a series of budget cuts in the late 2010s left a large community of researchers struggling to maintain publication output under a competitive funding scenario (Quintans-Júnior et al., 2020). All of these issues raise concerns about the replicability of published results – something that has not been evaluated systematically in the country.

With this in mind, we set up the Brazilian Reproducibility Initiative, a multicenter assessment of the replicability of experiments published by Brazilian institutions using common laboratory methods (Amaral et al., 2019; Neves et al., 2020). Our assessment of a representative sample of biomedical experiments published by national researchers aims to provide systematic data to raise awareness of the issue and inform scientific policy. In addition to generating data on replicability, the project’s first-person, naturalistic approach can identify challenges faced by the laboratories conducting replications, which are drawn from institutions similar to those that produced the original results. Consequently, our findings can offer insights into problems and potential solutions that are relevant to the country’s reality, as well as guide future replication efforts in lab biology.

## Methods

### Lab recruitment

The general rationale for the project has been described previously (Amaral et al., 2019). We began by systematically reviewing a random sample of life sciences articles from Brazil to identify common experimental models and methods (see https://osf.io/f2a6y). Based on these findings, we chose 10 experimental methods, using rodents or cell lines as models, to select studies for replication (see https://osf.io/qhyae). We then opened a public call for Brazilian labs that could replicate experiments using these methods and models, advertised by email, social media and lectures in conferences and institutions, to which 73 labs initially responded. Based on the expertise of respondents and a feasibility analysis by the coordinating team, we selected 3 outcome assessment methods for replication: the MTT (3-[4,5-dimethylthiazol-2-yl]-2,5 diphenyl tetrazolium bromide) reduction assay, the reverse transcription polymerase chain reaction (RT-PCR) and the elevated plus maze test of anxiety (EPM; see https://osf.io/qxdjt for details). Three further calls for labs were opened later to fill in specific gaps, leading to another 24 responses.

### Experiment selection

We searched the Web of Science for publications from 1998 to 2017 with affiliations in Brazil, using R code to identify the methods listed above (see https://osf.io/57f8s for details and https://osf.io/4sy2g for code). We then manually screened for articles that met the following criteria:

(a) had at least 50% of authors (including the corresponding one) affiliated with a Brazilian institution;

(b) had at least one quantitative result using one of the selected methods and models that (c) was statistically significant, (d) was mentioned in the title or abstract and (e) used only commercially available materials (for details, see https://osf.io/u5zdq). A data extraction step collected information on the biological models, experimental procedures and results for the first comparison in the article that presented a statistically significant difference between two groups. Manuals used for extraction were based on existing guidelines for methods reporting (Bustin et al., 2009; Kilkenny et al., 2010; NPQIP Collaborative group, 2019), and are available at https://osf.io/udjr7, https://osf.io/tr6xa, https://osf.io/7uhb6 and https://osf.io/rkvtm. After this step, we excluded experiments that did not report standard deviation, standard error of mean or confidence interval, or had an estimated cost greater than BRL 5,000 (around USD 1,300 at the time) per replication.

We then shared summaries of the eligible experiments’ methods with collaborating laboratories, according to their expertise, to confirm their capability to replicate them. Our replication sample comprised the first 20 experiments with each method that could be assigned to three independent labs. If labs withdrew from the project before completing their replications, those experiments were reassigned to other labs within our network or to new labs recruited by additional calls. Further information about (a) the selected experiments, (b) the original article, journal, authors and respective institutions, and (c) the teams performing replications were collected as outlined in https://osf.io/enjxy.

### Protocol development

For each experiment, the coordinating team transcribed the methodological information available in the original article into a protocol, with missing details left as gaps to be filled. Each lab assigned to replicate an experiment received this protocol, without information about the original publication or results (for details, see https://osf.io/gsvy2). They were instructed to fill in protocol gaps and elaborate on procedures when necessary, keeping as close to the original methods as possible (see instructions at https://osf.io/29vh7). Adaptations were allowed in cases of necessity (e.g. different equipment or unavailable reagents), or when the replicating lab considered the original protocol inadequate to measure the desired outcome, either due to methodological errors (e.g. incorrect primer sequences) or to significant risk of bias (e.g. lack of a vehicle group). Guidelines used to decide about adaptations are available at https://osf.io/e7zs9.

Protocols then received at least two external revisions, usually by another lab in the network that was not performing the same experiment and by a member of the coordinating team, both of whom had access to the original publication (see https://osf.io/g6ph5). Instructions to reviewers are available at https://osf.io/k97r4. For some protocols, a second coordinating team member replaced the independent lab as a reviewer. Reviewer comments were incorporated into the protocol as additional questions for revision and completion by the replicating labs. This process was iterated until the lab and the coordinating team deemed the protocol complete. Experiments involving animals were submitted to local institutional animal ethics review boards at this stage, and protocols were adjusted if required. After approval, the final version was reviewed once more by an independent member of the coordinating team, who raised additional questions if necessary, and then preregistered at the OSF (see https://osf.io/vzam6/ for the complete version history of registered protocols). A flowchart of the protocol development process is presented in **Figure S1**.

Sample size calculations were performed to achieve 95% statistical power to detect the original standardized mean difference in each individual replication (see details at https://osf.io/wxzr7). Code and data for the calculations are available at https://osf.io/vs5rp and https://osf.io/atkd7, respectively.

Authors of the original articles were contacted for missing information and raw data (see contact protocol at https://osf.io/3dn4x), but these were intended for future analyses and were not used in developing the replication protocols.

### Researcher predictions

Before the start of experiments, an open call was issued to experimental researchers willing to make predictions regarding the success of individual replications. Seventy participants selected one of the three methods and received a survey containing a summary of the 20 experiments associated with that method, including the original result and associated statistics. They were asked to predict (a) the probability of replication success, (b) the expected effect size in the replication, and (c) how technically challenging the replication would be (see details at https://osf.io/tm76h and full survey at https://osf.io/29mq5). Fifty-seven of these researchers also participated in prediction markets, in which they received USD 50 in vouchers to bet on the replication success of individual experiments. An analysis plan for this part of the project, which will be explored in future publications, was preregistered at https://osf.io/pjhgd, and anonymized survey data is deposited at https://doi.org/10.7910/DVN/2RLSMG.

### Experimental procedures

For each experiment, a custom data collection spreadsheet was built by the coordinating team based on the protocol. This was meant to document not only the experimental results but also the execution of each protocol step to make the experiment auditable. Labs received these spreadsheets along with a detailed manual on how to execute and document the replications (https://osf.io/ackyd) and on how to deal with protocol adaptations when necessary (https://osf.io/e7zs9).

Experimental materials were acquired by the coordinating team and delivered to the lab performing the replication, unless they were already available there. Labs were expected to only use materials within their expiration date; however, due to the frequent postponing of experiments because of COVID-19 restrictions or difficulties with suppliers, the use of reagents with expired validity was allowed if their activity could be clearly demonstrated – either by the method demonstrably working as expected (e.g. PCR kits, molecular weight scales) or by a separate test of biological activity with a prespecified expected result (e.g. a test of cell growth with a particular culture medium). These cases were documented in data collection forms, compiled by the coordinating team and evaluated by the validation committee (see below).

Labs then performed replications and sent the data collection sheets to the coordinating team when experiments were completed. If any difficulties arose, labs were free to contact the coordinating team for orientations. If results suggested a failure to implement the method adequately (Neves & Amaral, 2020), labs were allowed to repeat experiments (see **Figure S1**), and general guidelines used to deal with these situations were later compiled in https://osf.io/m35s6.

### Post-experiment debriefing

After an experiment was finished, data collection spreadsheets were sent to the coordinating team, who reviewed them and created a document with questions concerning (a) experimental steps deviating from the preregistered protocol and (b) unclear or missing information. These were sent to the replicating lab, which was asked to confirm the observations on protocol deviations, answer the coordinating team’s questions and rate (on a scale of 1 to 5) how much the executed protocol deviated from the preregistered one (data available at https://osf.io/g9jsp, https://osf.io/64whp and https://osf.io/dfp3h).

After undergoing this step and answering further questions by the coordinating team if needed, labs received access to the original article from which the experiment was drawn. They were asked to fill in a form answering whether they felt the original result was successfully replicated, with a justification for their answer. They also rated how relevant the differences between the original protocol and the replication were (on a scale of 1 to 5). Subjective assessment decisions were reviewed by e-mail with the labs if (a) justifications suggested answers concerned the reproducibility of methods rather than results or (b) errors or inconsistencies in replication results were detected and corrected in subsequent steps. Data is available at https://osf.io/dqwr4, and more details on each debriefing step are presented at https://osf.io/xgth2.

Based on the responses to the first debriefing form, a final notes document on each experiment was compiled by the coordinating team, containing (a) a list of deviations from the original protocol, (b) a list of additional observations on the experiment, (c) a list of reagents used after the expiration date (if any) and (d) clarifications about the data collection spreadsheet.

### Validation of experiments

To adjudicate whether the performed experiments constituted valid replications and should be included in the main analysis, a validation committee was formed for each method, including the coordinating team and members of participating labs that used that method. Each experiment was evaluated by three members of this committee, who received the registered protocol, the original article, the final notes document, and (in some cases) the data collection spreadsheets and/or additional material, but with no access to the replication results. Each member of the validation committee received instructions (https://osf.io/rnemd) on how to evaluate these documents and was asked to answer how much the executed experiment deviated from the registered protocol (on a scale of 1 to 5) and whether they considered it a valid replication, providing justifications for their answers.

If any of the three evaluators or the replicating lab considered the replication invalid or attributed a deviation rating greater than 3, or if the sum of the 4 evaluation scores was greater than 10, the experiment was discussed by the validation committee to decide whether it should be included in the primary analysis. The committee could also suggest the exclusion of individual experimental units, or the use of an experimental unit different from the one suggested by the lab. Committee members did not participate in the discussion of their own replications. After all experiments were discussed, decisions were reviewed for consistency, and inconsistent decisions were rediscussed to align criteria. Moreover, if any additional issues were raised during the analysis or qualitative assessment steps (see below), experiments were rediscussed by e-mail with the validation committee and decisions were changed if necessary. **Figure S2** provides an overview of the debriefing and validation process; for more details, see https://osf.io/e3fjg. Decisions are available at https://osf.io/9ta45.

### Lab self-assessment, project evaluation and qualitative assessment of replications

After the conclusion of the validation process, replicating labs received the final notes documents, validation committee scores and validation decisions. They were then asked to fill in a final debriefing form (see https://osf.io/xgth2) where they should answer about their agreement with these scores and decisions. They were also asked to elaborate on reasons for protocol deviations, whether those could have been prevented, and what they would change in the replication protocol if they were to start over. Data for this step is available at https://osf.io/dp54y.

In addition to these lab responses, individual lab members were asked to optionally fill in an anonymous project evaluation form, in which they could evaluate the project and their participation, and elaborate on challenges, difficulties, and learning opportunities. The 121 responses to this form (https://osf.io/gdvam) were complemented by 18 semi-structured interviews with project participants. Responses to open questions about protocol deviations and project difficulties were categorized by the coordinating team following a taxonomy available at https://osf.io/5gjb7. After this, the coordinating team built its own list of difficulties faced by the project. For more on the project evaluation process, see https://osf.io/nfr6y.

Finally, each experiment’s results were sent to the labs that replicated it, along with the original article, information provided by the original authors (when available), replication protocols and validation decisions. Labs were asked to provide an assessment of the aggregate outcome of the replication, and their responses were discussed in a live meeting involving participants from the involved labs and the replication team. Based on these discussions, the coordinating team prepared qualitative assessments of each replication, which will be analyzed in future publications. More on this step can be found at https://osf.io/w5z9a.

### Data cleaning and checking

Primary data from the experiments recorded in data collection spreadsheets (optical density measurements for MTT and conventional PCR, Ct values for quantitative PCR (RT-qPCR), and behavioral outcomes for EPM) were manually compiled by the coordinating team into a single spreadsheet for each method, along with experiment-level metadata (available at https://osf.io/pwfze). When inconsistencies in calculations or discrepancies between subsections of the data collection spreadsheets were detected, they were cleared up by written contact with the replicating lab. After this, results were summarized by code for each experiment into spreadsheets and scatter plots and shared with the labs, who were asked to check for discrepancies with their calculations. If issues were found, the lab and coordinating team reviewed them via email until the sources of errors were identified and corrected (see https://osf.io/58vsx for details).

Nevertheless, during online discussions for qualitative assessment of replications – which occurred after publication of the initial version of the preprint – we found that several errors in data compilation were not detected by this checking process. The coordinating team then proceeded to check each experiments’ data collection sheet individually against the compiled results and used the qualitative assessment meetings to review inconsistencies with the labs. This led to revisions of data in 26 replications (18% of the total), of validation decisions in 7 replications (5%) and of subjective assessments of results in 6 replications (4%), as well as adjustments in documentation for individual experiments. A complete list of changes is available at https://osf.io/4tu2v.

## Data analysis

The protocol for analyzing replication rates and predictors was initially registered at https://osf.io/9rnuj, before any experimental results had been reviewed by the coordinating team. It was updated at a time point when experimental results were available, but no analyses had been performed, to account for steps added after the experiments (such as the validation process). The final dataset used for analysis is deposited at https://doi.org/10.7910/DVN/ZJRDIV. Analysis code in R was developed based on this protocol and is available both in the data repository and at https://github.com/BrazilianReproducibilityInitiative/bri-analysis. A list of deviations, additions and specifications added to the protocol after the analysis had started is available at https://osf.io/9hj7t.

In summary, results from the available replications of each experiment were aggregated by meta-analysis using the R package metafor (Viechtbauer, 2010), with the log-transformed ratio of means as the effect measure for MTT and EPM. For RT-qPCR experiments, we used the mean difference in ΔCt values (which is already in log scale), while for conventional PCR we used the log-transformed relative band density between the gene of interest and the reference gene. The results of these meta-analyses were used for comparison with the original result (also transformed into the natural logarithm of the ratio of means). Paired normalization between units in the experimental and control groups was implemented for all MTT experiments, as absolute optical density values were not considered commensurable across experimental units measured in different days. For PCR experiments, pairing was used only when this was the case in the original study, as described in the methods or inferred from the lack of error bars in the control group (see https://osf.io/wxzr7).

Replication rates for the sample were calculated on the basis of 5 dichotomous criteria: (a) whether the original estimate was within the 95% prediction interval of a random-effects meta-analysis of the replication, (b) whether the estimate of a random-effects meta-analysis of the replications was within the 95% confidence interval of the original results, (c) whether the point estimate of a fixed-effects meta-analysis of the replication was statistically significant at p < 0.05 and had the same sign as the original result, (d) whether at least half of individual replications were statistically significant in the same direction as the original result and (e) whether at least half of labs considered the original result successfully replicated in their subjective assessment. Prediction and confidence intervals were based on t distributions with degrees of freedom based on the number of experimental units, combined between groups within replications and between replications within meta-analyses using the Welch-Satterthwaite equation (Welch, 1947). Sensitivity analyses use (a) z distributions, (b) the Knapp-Hartung approach to calculate degrees of freedom (Knapp & Hartung, 2003) or (c) a single mean of the experimental units from all replications rather than meta-analysis (with data for each experimental unit normalized by their respective control unit in paired experiments or by the control mean of the respective replication in unpaired ones). Agreement between replication criteria was calculated by Cohen’s kappa for pairs and Fleiss’ kappa for the aggregate of the 5 criteria.

These rates were initially calculated for the primary analysis set, which included only the replications endorsed by the validation committee. Rates were also calculated for other sets of replications, namely (a) all replications, (b) all replications considered valid by the lab, (c) only experiments with at least 2 valid replications, (d) only experiments with at least 3 valid replications, (e) only replications achieving 80% post-hoc power on aggregate to detect the original relative difference in means, considering the standard deviation, sample size and correlation between pairs found in the replication (see https://osf.io/tbkvz for experiments within each analysis set and results of post-hoc power calculations). The primary analysis used the experimental unit as defined by the validation committee, while sets (a) and (b) above considered the lab’s definition. A multiverse analysis using all possible replication criteria, sets of experiments and analysis decisions is presented as a specification curve (Simonsohn et al., 2020).

The correlation between effect sizes from the replication and original experiments was evaluated using Pearson’s *r*, with sensitivity analyses removing a prominent outlier or using Spearman’s ρ. Coefficients of variation from the original study were compared to the mean coefficient of variation of its replications using Wilcoxon’s signed rank test. The mean absolute difference between the effect sizes of the original study and its replications (both expressed as log ratios of means) was compared to that between individual replications of the same experiment, also using Wilcoxon’s signed rank test. These analyses were added after data collection and should thus be considered exploratory. Predictors of replication success at the level of original experiments and at the level of replications were evaluated using Spearman’s correlation coefficient as planned in the protocol. A complete list of tested predictors is available in the analysis protocol and list of deviations.

## Results

### Lab recruitment and selection of experiments

Throughout the project, 97 labs across Brazil applied to replicate experiments in response to 4 public calls. Of these, 75 joined the Initiative at some point and 56 contributed data. The geographic distribution of these labs is displayed in **Figure S3** and **Table S1**, with information on team members available in **Table S2**.

The selection of experiments for replication and their execution are summarized in **Figure 1**. A total of 317 experiments using the 3 selected methods (MTT, PCR and EPM) were identified by full-text screening and had protocol data extracted. 173 protocol summaries (https://osf.io/7kagj) were shared with participating labs until we could assign 20 experiments with each method to at least 3 labs. Information on selected experiments and their articles of origin can be found at https://osf.io/cyjsz and is summarized in **Table S3**, while replication protocols developed by labs are available at https://osf.io/vzam6/.

**Figure 1.**
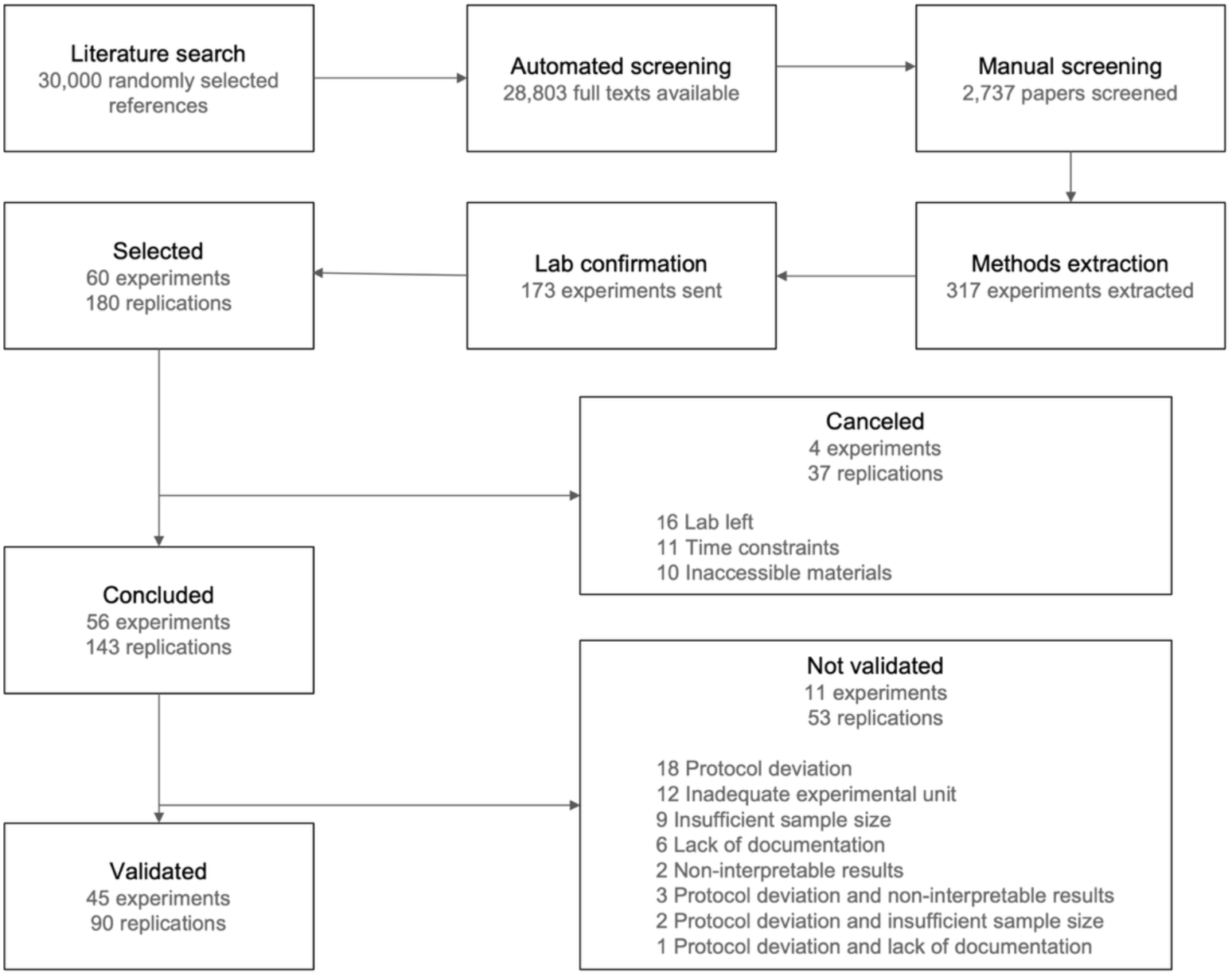
Flowchart describing the selection and execution of replication experiments. Top row shows the number of articles that underwent automated and manual screening. The methods of selected experiments were extracted, and their summaries were sent to labs to determine the final replication sample. Numbers of concluded and validated experiments (i.e. those with at least one replication) and replications are shown in the bottom rows, with reasons for lack of completion or invalidation shown on the right.

### Concluded experiments

Out of the 180 planned replications, 143 were carried out by labs, covering a total of 56 experiments, of which 34 had three replications, 19 had two replications and 3 had only one (**Table S4**). 136 of these had enough data (i.e. more than one valid data point in each group) to be used in meta-analyses. Canceled replications were due to inaccessible materials or to labs leaving the project or not finishing replications by the project’s deadline.

Experiments submitted by labs then underwent validation by our designated committee. Out of 143 replications, 53 were excluded in this step due to protocol deviations (18), using an inadequate experimental unit (12), insufficient sample size (9), insufficient documentation (6), non-interpretable results (2) or a combination of these reasons (6) (see **Figure 1**). Results of the validation process are available at https://osf.io/9ta45/ and a description of reasons for invalidation can be found in **Table S5**. Validation decisions for different interpretations of the experimental unit in cell culture experiments are detailed in **Table S6**. Labs agreed with the committee’s decisions on all validated experiments and on 77% of invalidated ones.

This left us with 90 valid independent replications covering 45 different experiments in our primary analysis. Five replications with insufficient sample size and n > 1/group were included in meta-analyses but not analyzed independently. Only 9 experiments remained with three valid replications, while 27 had two and 9 had only one. A list of materials concerning each replication is compiled at https://osf.io/6av7k/wiki/Individual_Replications/, while aggregated data for each analysis set is available at https://osf.io/6av7k/wiki/Aggregated_Replications/. For the primary analysis, unit-level data are available as tables and scatter plots at https://osf.io/uzm97/files/, and forest plots are available at https://osf.io/sx9gv. Qualitative assessments of experiments are available at https://osf.io/p7b9z.

### Replication rates

Replication rates for the 45 experiments included in the primary analysis are shown in **Table 1**. In 17 out of 39 experiments with multiple replications (44%), the original estimate was within the 95% prediction interval of the replication meta-analysis, while 13 out of 45 aggregate replication estimates (29%) and 23 out of 90 individual replications (26%) were within the original experiment’s 95% confidence interval. Nine (20%) aggregate estimates and 17 (19%) individual replications showed a statistically significant effect in the same direction as the original, with half or more of the available replications significant in 10 experiments (22%). Twenty-nine individual replications (32%) were considered successful by the replicating lab, with at least half of labs making this judgment in 19 experiments (42%). That said, many labs used loose criteria for success, such as an effect in the same direction or of comparable size, regardless of statistical analyses (see **Table S7**). Twelve (27%) experiments were replicated by at least half of the applicable criteria, and agreement between different criteria is shown in **Figure S4**, with Fleiss’ kappa coefficients of 0.54 for the 5 experiment-level criteria and 0.38 for the 3 replication-level criteria.

**Table 1.** Replication rates in the primary analysis. Replication rates for the primary analysis using multiple criteria. Effect size comparisons are based on random-effects meta-analyses, while same-sign significance is based on fixed-effects meta-analysis estimates. The 95% prediction interval criterion only uses experiments with more than one replication and thus has a different sample size. All statistical tests use t distributions based on the number of experimental units. PI, prediction interval; CI, confidence interval; MTT, 3-[4,5-dimethylthiazol-2-yl]-2,5 diphenyl tetrazolium bromide assay; PCR, reverse transcription polymerase chain reaction; EPM, elevated plus maze. For more information on replication criteria, see https://osf.io/9rnuj.

| By experiment | All | MTT | PCR | EPM |
| --- | --- | --- | --- | --- |
| Original in replication's 95% PI | **17/39 (44%)** | 8/15 (53%) | 8/13 (62%) | 1/11 (9%) |
| Replication in original 95% CI | **13/45 (29%)** | 5/17 (29%) | 5/14 (36%) | 3/14 (21%) |
| Same-sign significance (p<0.05) | **9/45 (20%)** | 4/17 (24%) | 3/14 (21%) | 2/14 (14%) |
| ≥50% replications significant | **10/45 (22%)** | 5/17 (29%) | 3/14 (21%) | 2/14 (14%) |
| ≥50% subjectively replicated | **19/45 (42%)** | 8/17 (47%) | 7/14 (50%) | 4/14 (29%) |
| By replication | All | MTT | PCR | EPM |
| Replication in original 95% CI | **23/90 (26%)** | 7/33 (21%) | 6/28 (21%) | 10/29 (34%) |
| Same-sign significance (p<0.05) | **17/90 (19%)** | 10/33 (30%) | 5/28 (18%) | 2/29 (7%) |
| Subjectively replicated | **29/90 (32%)** | 12/33 (36%) | 11/28 (39%) | 6/29 (21%) |

A major caveat for the interpretation of these replication rates is the low statistical power of some replications. Although we calculated sample sizes for each individual replication to have 95% power to detect the original standardized difference (with the primary analysis including only those achieving at least 80% power for this purpose), coefficients of variation were on average much higher among replications, particularly for PCR experiments (**Figure S5**). This led statistical power to detect the original relative difference to be lower than planned for many experiments (**Figure S6**), biasing replication rates based on statistical significance downwards and those based on prediction intervals upwards (as also occurs when there is low agreement among replication results). To address this, we conducted an exploratory *a posteriori* power analysis considering only experiments with at least 80% statistical power for the aggregate of replications (n=35), using the original relative difference and the variability achieved in replications. This leads to an increase in replication rates based on statistical significance and a decrease in those based on prediction intervals, with 38% of original effects within the replications’ 95% prediction interval, 31% of replication estimates within the original 95% confidence interval and 26% of experiments statistically significant in the same direction as the original (**Table S8**).

**Table 2** displays replication rates for these and other sets of experiments, including all experiments irrespectively of the validation results (n=56), all experiments considered valid by labs (n=56), and only experiments with at least two (n=36) or three (n=9) valid replications (broken down by method in **Tables S9-S12**). Disregarding the validation process increases sample size but has little impact on replication rates, whereas analyzing only experiments with multiple replications leads to higher rates by some criteria, but with a lower sample size. Replacing t by z distributions in meta-analyses (**Tables S13-S14)** decreases replication rates based on confidence intervals, which become narrower, but increases significance rates. Basing degrees of freedom for meta-analyses on the Knapp-Hartung approach (**Tables S15-S16**) increases replication rates based on prediction intervals but drastically lowers those based on meta-analytical significance, as estimates incorporate more uncertainty. Finally, synthesizing replications by a mean of all experimental units rather than by meta-analysis (**Tables S17-18)** lowers replication rates, as the precision of aggregate estimates is decreased by individual replications with high variability.

**Table 2.** Replication rates for different analysis sets. Replication rates for the primary and secondary analyses. Effect size comparisons are based on random-effects meta-analysis, while same-sign significance is based on a fixed meta-analysis estimate. The 95% prediction interval criterion only uses experiments with more than one replication and thus has a different sample size. Subsets for secondary analyses include all experiments judged valid by the replicating lab (Lab’s Choice), all concluded experiments (All Exps) – both of which use the experimental unit as defined by the lab rather than by the validation committee –, only experiments with at least 2 (≥ 2 Reps) and 3 replications (3 Reps), and only experiments with ≥ 80% *a posteriori* power using the original relative difference and the variability achieved in replications. All statistical tests use t distributions based on the number of experimental units. PI, prediction interval; CI, confidence interval. For more information on replication criteria, see https://osf.io/9rnuj.

| By experiment | Primary | Lab's choice | All Exps | ≥ 2 copies | 3 copies | ≥80% power |
| --- | --- | --- | --- | --- | --- | --- |
| Original in replication's 95% PI | **17/39 (44%)** | 22/43 (51%) | 25/52 (48%) | 16/36 (44%) | 4/9 (44%) | 13/34 (38%) |
| Replication in original 95% CI | **13/45 (29%)** | 15/56 (27%) | 14/56 (25%) | 11/36 (31%) | 4/9 (44%) | 11/35 (31%) |
| Same-sign significance (p<0.05) | **9/45 (20%)** | 13/56 (23%) | 12/56 (21%) | 9/36 (25%) | 2/9 (22%) | 9/35 (26%) |
| ≥50% replications significant | **10/45 (22%)** | 11/56 (20%) | 10/56 (18%) | 10/36 (28%) | 1/9 (11%) | 9/35 (26%) |
| ≥50% subjectively replicated | **19/45 (42%)** | 19/56 (34%) | 19/56 (34%) | 15/36 (42%) | 3/9 (33%) | 12/35 (34%) |
| By replication | Primary | Lab's choice | All Exps | ≥ 2 copies | 3 copies | ≥80% power |
| Replication in original 95% CI | **23/90 (26%)** | 31/116 (27%) | 35/136 (26%) | 21/81 (26%) | 11/27 (41%) | 21/76 (28%) |
| Same-sign significance (p<0.05) | **17/90 (19%)** | 22/116 (19%) | 23/136 (17%) | 17/81 (21%) | 4/27 (15%) | 16/76 (21%) |
| Subjectively replicated | **29/90 (32%)** | 35/116 (30%) | 41/136 (30%) | 25/81 (31%) | 9/27 (33%) | 21/76 (28%) |

We also evaluated the impact of analysis decisions made for specific methods after the protocol was registered. Pairing MTT experiments only when the original was presumably paired markedly increases coefficients of variation, decreasing statistical power and replication rates based on statistical significance (**Table S19**). Analyzing relative expression in a linear scale for PCR experiments (**Table S20**) also leads to increased variability, lower rates of statistical significance and wider prediction intervals. A multiverse analysis for all combinations of experimental sets, replication criteria and analysis options is presented as a specification curve in **Figure 2** and shows a median replication rate of 25% (interquartile range = 19-33%).

**Figure 2.**
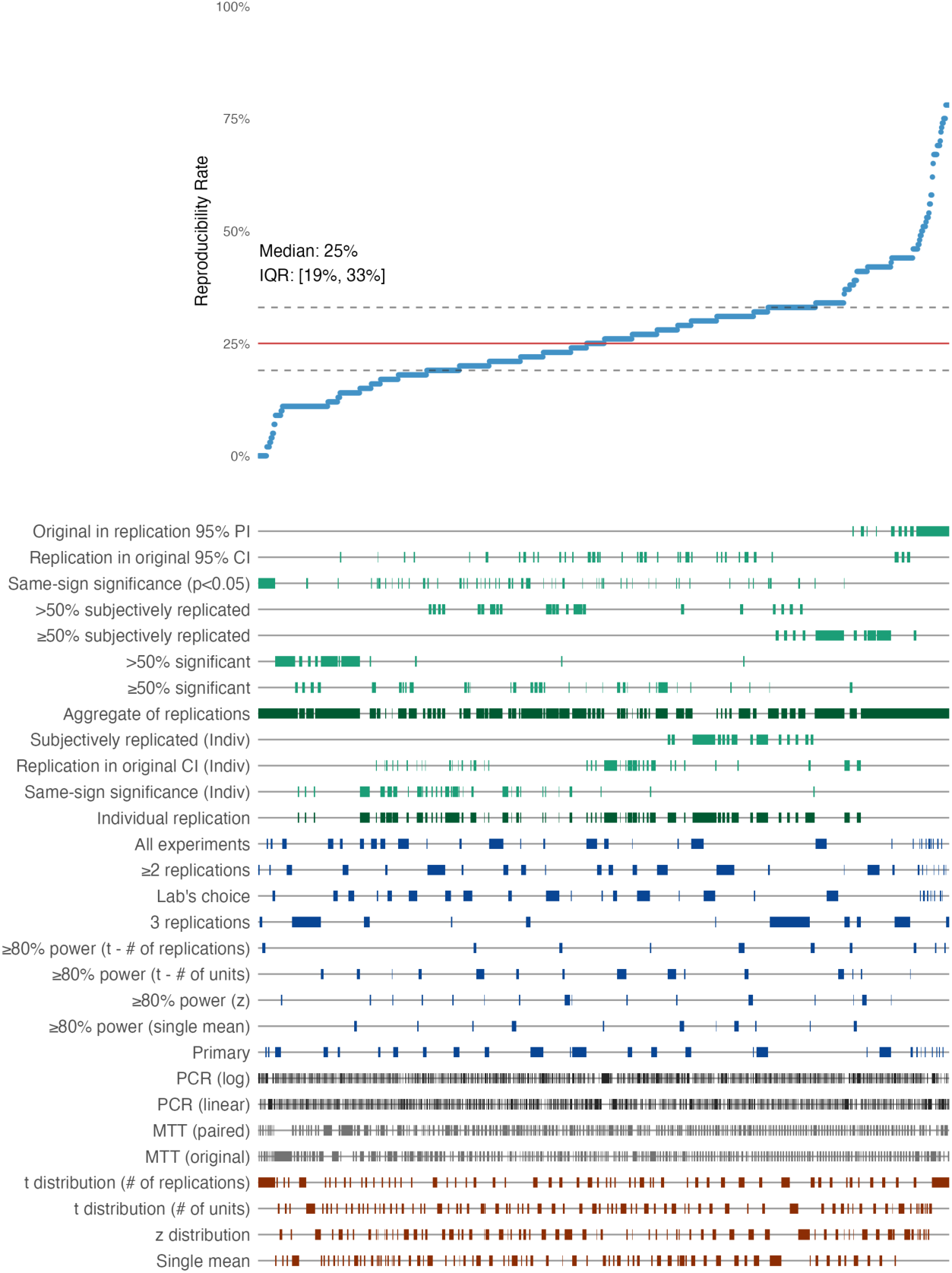
Specification curve analysis of replication rates. Curve shows replication rates arising from the combination of 10 replication criteria (green; 5 criteria for the aggregate of experiments, with 2 different ways to resolve ties for those involving voting, and 3 criteria for individual replications), 8 sets of experiments with different criteria for inclusion (blue), 2 ways of handling PCR (black) and MTT (grey) data, and 4 different approaches for statistical analysis (red). Replication rates range from 0 to 78%, with a median of 25% and an interquartile range (IQR) of 19 to 33%.

### Effect size and variability comparison

A comparison between the effect sizes from each original experiment and those of replications in the primary analysis is shown in **Figure 3A**, while the correlation between the aggregate effect size of replications and the original one is shown in **Figure 3B**. The high linear correlation (r = 0.82, p = 6.4×10^-12^) is due to one prominent outlier with very large effect sizes in both the original and replication; removal of this experiment leaves a weak correlation with r = 0.22 (p = 0.16; **Figure 3C**). Using a non-parametric approach for the whole sample also leads to a lower correlation coefficient (Spearman’s ρ = 0.35, p = 0.02).

**Figure 3.**
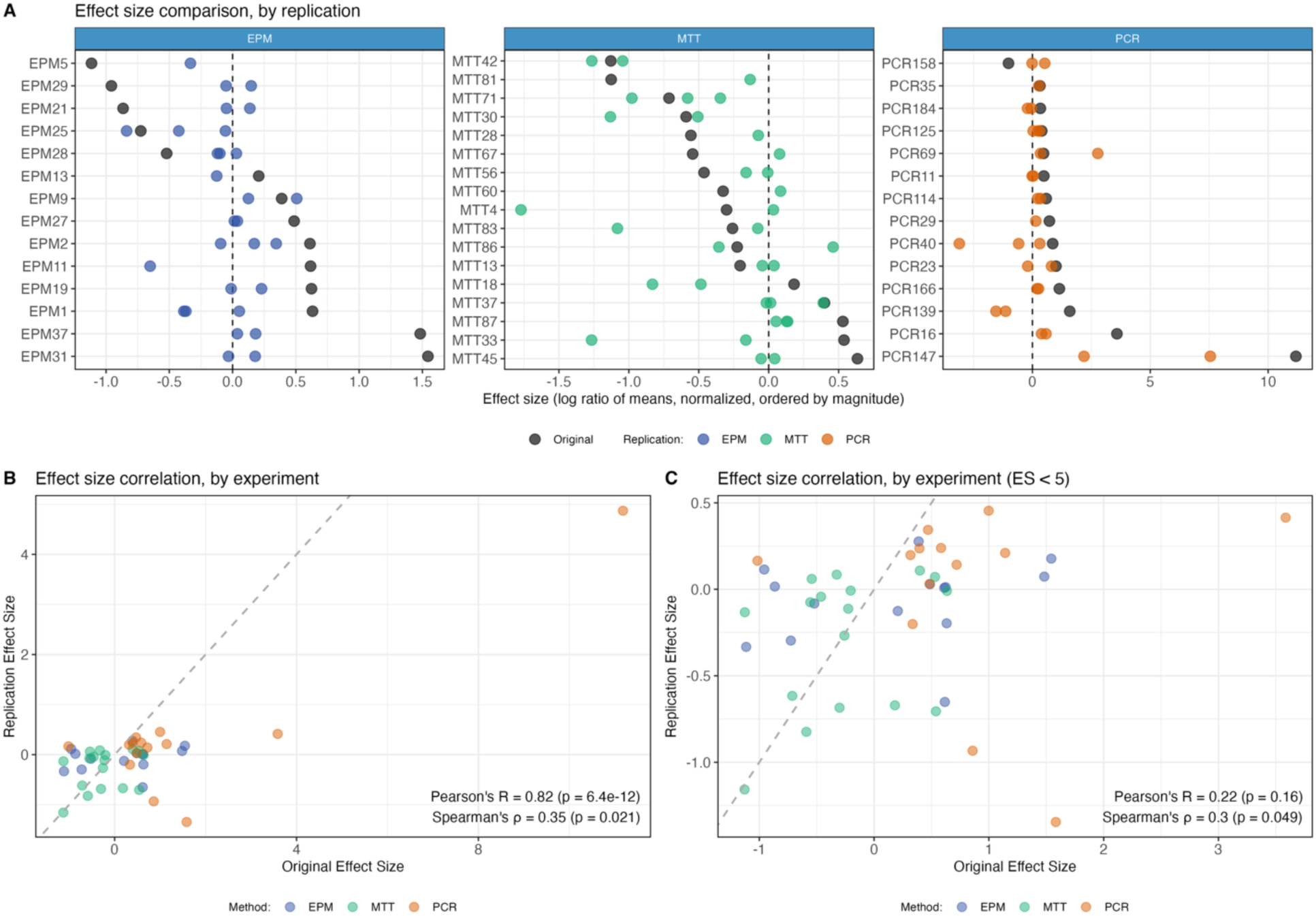
Effect size comparisons and correlations. **(A)** Effect size comparison between original experiments (dark) and individual replications (light) for EPM (blue), MTT (green) and PCR (orange) experiments, ordered by the original effect size. X axes represent effect sizes as the natural logarithm of the ratio between the means of the experimental and control groups in behavioral outcomes (EPM), optical density (MTT) or relative gene expression (PCR), with 0 indicating no difference between groups. **(B)** Correlation between the effect sizes of original experiments (x axis) and the aggregate result of their respective replications (y axis), again expressed as log ratios of means (r = 0.82, p = 6.4×10^-12^; ρ = 0.35, p = 0.02). Colors are the same as in A. Dashed line indicates equivalent effect sizes between original and replication. **(C)** The same analysis excluding the prominent outlier in PCR147, yielding a much weaker linear correlation (r = 0.22, p = 0.16; ρ = 0.3, p = 0.05). All results use the primary analysis set and t distributions based on the number of experimental units.

41 out of 45 effect sizes (91%) were smaller in the replication aggregate than in the original, with a median ratio between the original and replication relative effect sizes (both expressed as ratios of means) of 1.71 (1.51 for MTT, 1.75 for PCR and 2.01 for EPM) (**Table 3**). This expresses a ratio between ratios, which is used for mathematical purposes to include effects in the opposite direction: median relative differences (80% in the original vs. 8% in the replication when considering effect direction) are 90% smaller in the replications. While this may reflect exaggeration of original effect sizes or publication bias (Ioannidis, 2008), some degree of inflation is expected even in its absence, as our selection process filtered for original experiments with significant differences. In an exploratory analysis, we found that coefficients of variation were lower in the original than in the average of replications in 33 out of 45 experiments (73%), with a median ratio of 0.55 (Wilcoxon’s signed rank test, p = 1.3×10^-5^). This difference was greater in PCR experiments (ratio of 0.28, p = 2.4×10^-4^) than in MTT and EPM ones (ratios of 0.85 and 0.74, p = 0.15 and 0.05, respectively). Sign errors (i.e. significant effects in the opposite direction of the original) were infrequent, occurring in 2 out of 45 experiments (4%) and accounting for 18% of significant replication results.

**Table 3.**
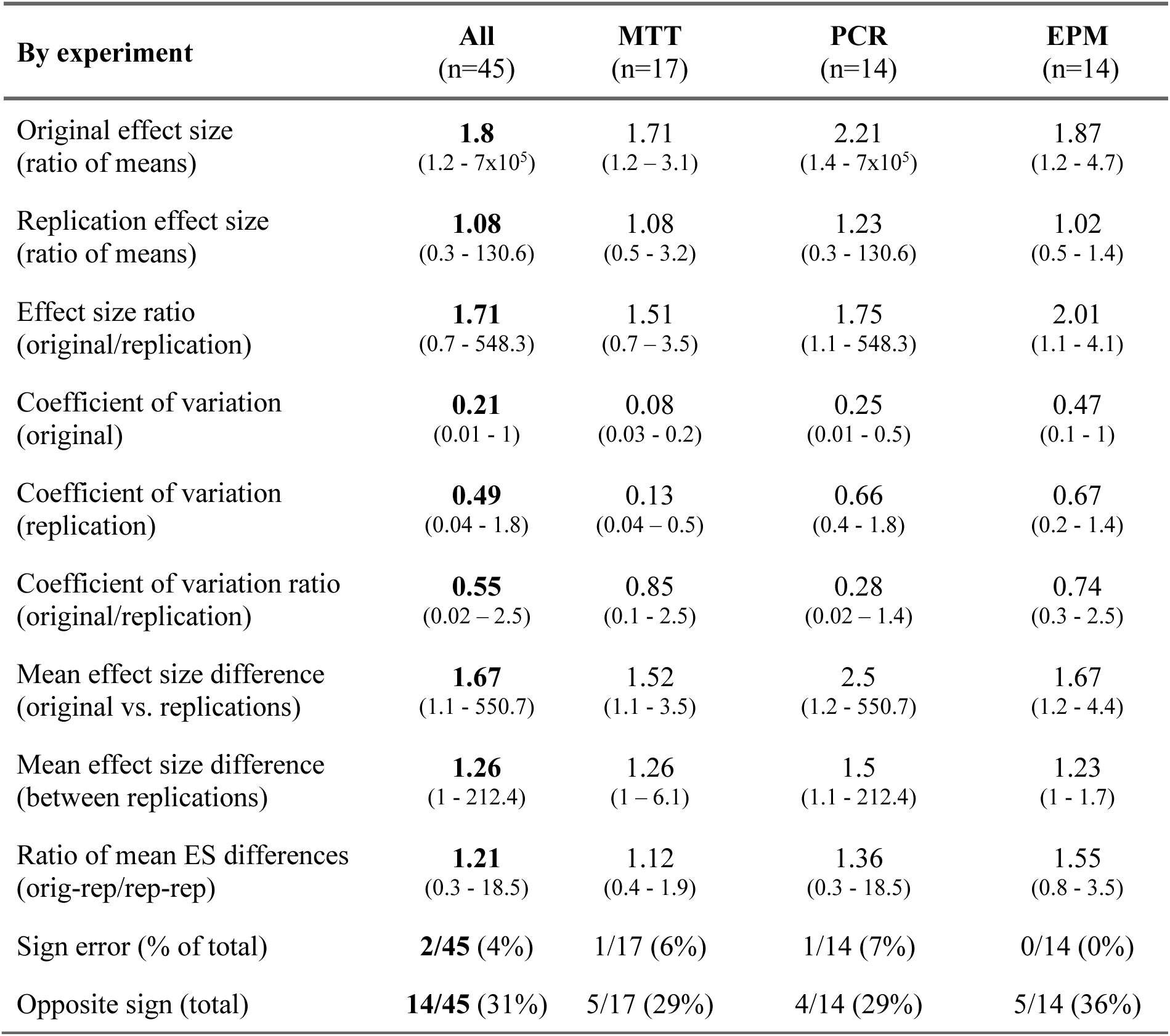
Comparisons between results of replications and original studies in the primary analysis. Continuous variables are shown as median (range), while categorical ones are shown as proportion (percentage). Original effect sizes represent the ratio between the higher and lower mean of the two groups in the original study and are thus always above 1. Replication results respect the same order as the original: therefore, effects above 1 are in the same direction, and those below 1 are in the opposite direction. Ratios between original and replication effect sizes are thus ratios between ratios and are calculated by exponentiating differences between log ratios. Coefficients of variation are calculated as the pooled mean of both groups divided by the pooled standard deviation; for PCR experiments, this is done for relative expression values in linear scale. Mean effect size differences are obtained from absolute differences between effect sizes in log ratio of means, in order to measure discrepancies between the original experiment and its replications or between individual replications. These are also exponentiated and thus represent the mean ratio between the higher and lower value. Sign errors refer to effects in the opposite direction of the original with p < 0.05, while opposite sign (total) includes all differences in the opposite direction (irrespective of significance). All results use the primary analysis set and t distributions based on the number of experimental units. ES, effect size; MTT, 3-[4,5-dimethylthiazol-2-yl]-2,5 diphenyl tetrazolium bromide assay; PCR, reverse transcription polymerase chain reaction; EPM, elevated plus maze.

Our multicenter design also allowed us to compare the variation between individual replications to that between replications and original studies in an exploratory analysis. Relative differences (expressed as ratios between the higher and lower mean) between original effects and individual replications were larger than those between replications of the same experiment, with a median ratio of 1.21 (Wilcoxon signed-rank test between pairs, p = 0.01). Differences were larger and more consistent among EPM studies (median ratio = 1.55, p = 0.007) than among PCR (ratio = 1.36, p = 0.19) or MTT experiments (ratio = 1.12, p = 0.64) (**Table 3**). This suggests that part of the irreplicability observed in our study is due to factors specific to published experiments that were not present in replications; nevertheless, a sizable amount of variation was still observed among replications, as can be seen in **Figure 3A**.

### Predictors of replication success

We analyzed whether multiple factors at the level of the original experiment and replication were correlated with replication success, as measured by the strength of evidence for an effect in the same direction (the t value for the replication or aggregate of replications) and by effect size exaggeration (the difference between the effect sizes of the original experiment and replication, both expressed as log ratios of means). A summary of these correlations in the primary analysis is shown as a heat map in **Figure 4**, while individual scatter plots are available at https://osf.io/fdpbe.

**Figure 4.**
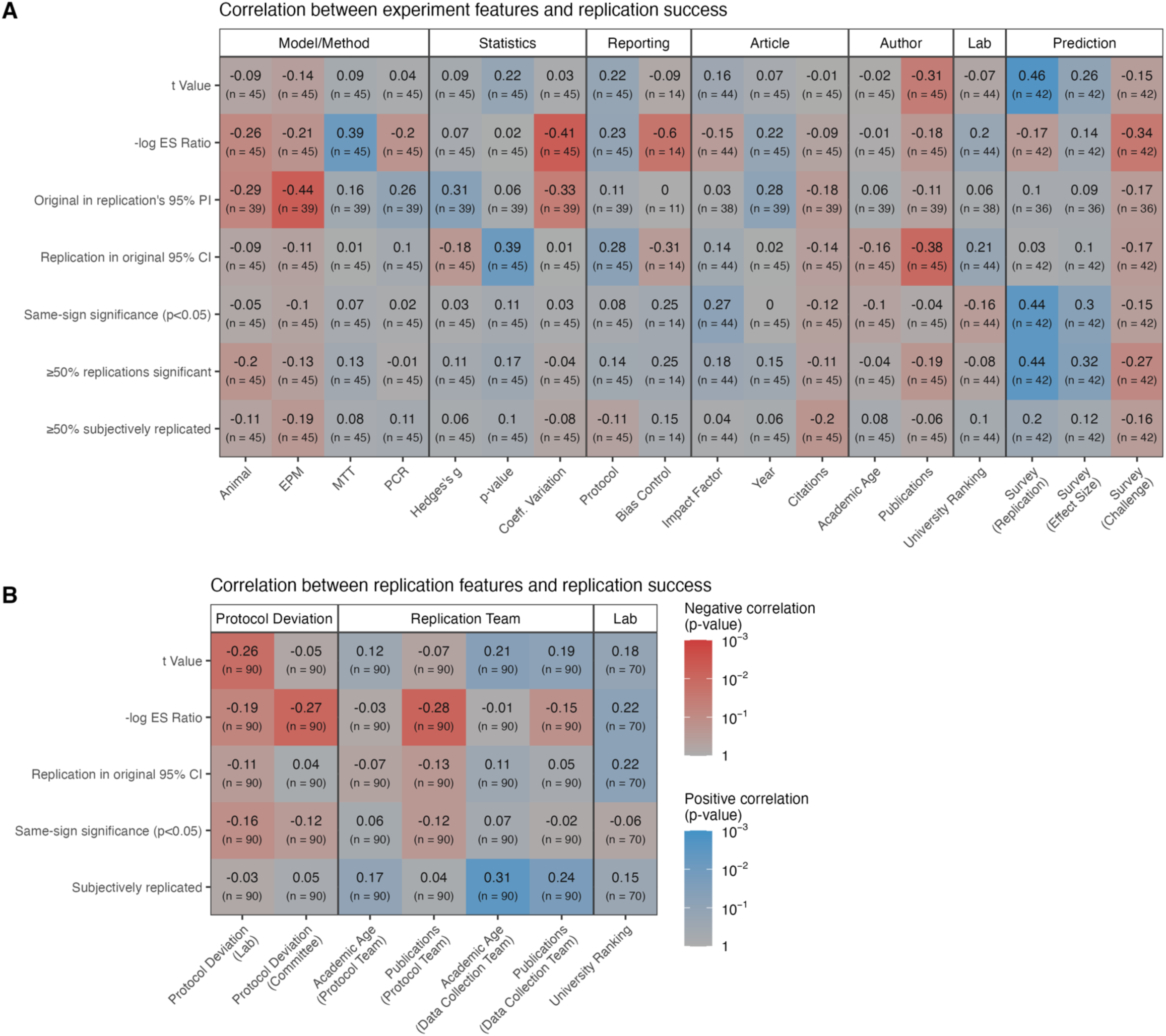
Predictors of replication success. **(A)** Predictors of replication success at the level of original experiments (x axis) in the primary analysis. Categories include experimental model and method (animal vs. cell, EPM/MTT/PCR vs. others), statistics (original standardized effect size, p value and coefficient of variation), reporting (protocol reporting score, bias control measures), article features (journal impact factor, year and citations in the first 2 years), author features (years since first publication and number of publications at the time of original article publication), lab features (university ranking) and researcher predictions (about replication probability, expected effect size and difficulty). For details on each predictor, see https://osf.io/9rnuj. Y axis represents different continuous (aggregate replication t value, effect size differences in log scale) and dichotomous outcomes (same as in Table 1); effect size differences are expressed as replication minus original (i.e. with a sign opposite to that presented in the text) so that blue indicates correlations with less exaggeration/higher replicability and red with more exaggeration/lower replicability. Numbers show univariate correlations as Spearman’s ρ, while color intensity represents p values. **(B)** Predictors at the level of replications, including degree of protocol deviation (judged by the lab and validation committee), features of the replication team (mean number of years since first publication and number of publications by the protocol and data collection teams) and of the lab (university ranking). Other conventions are the same as in A, but outcomes and sample size refer to individual replications. Scatter plots for individual correlations are available at https://osf.io/fdpbe.

At the original experiment level (**Figure 4A)**, replication success as measured by t value was mainly correlated with researcher predictions about the probability of replication success (ρ = 0.46, p = 0.002) (see **Table S21** for data on survey participants and **Figure S7** for survey results). Predictions were also correlated with dichotomous same-sign significance (ρ = 0.44, p = 0.004), which was the outcome participants were directly asked to predict. Conversely, t values were negatively correlated with the number of publications by the original article’s last author in the 5 years preceding the replicated publication (ρ =-0.31, p = 0.04). Effect size exaggeration was greater for experiments with higher coefficients of variation in the original experiment (ρ = 0.41, p = 0.005) – an expected result when original results are filtered for significance –, and lower for MTT (ρ =-0.39, p = 0.008), and cell line experiments (ρ =-0.26, p = 0.08), although these variables are heavily correlated among themselves, making confounding likely (see **Figure S8** for correlations among predictors). Exaggeration was also correlated with researcher predictions on how challenging the replication would be to perform (ρ = 0.34, p = 0.03), worse reporting of the experiment’s methods (ρ =-0.23, p = 0.13) and, surprisingly, with higher use of bias control measures in EPM experiments (ρ = 0.6, p = 0.02), although the latter analysis had a very low sample size (n=14). Most of these correlations are weak and, due to the multiplicity of tested predictors, could plausibly have arisen by chance.

In an exploratory analysis at the replication level (**Figure 4B**), t values were negatively correlated with the degree of change between the original article and replication, as assessed by the replicating lab (ρ =-0.26, p = 0.01); however, as this score was attributed after seeing the results, unsuccessful replications could have led researchers to attribute greater value to protocol changes. Effect size exaggeration was positively correlated with this score (ρ = 0.19, p = 0.07), as well as with that attributed by the validation committee, which was blind to the replication results (ρ = 0.27, p = 0.009). More experienced data collection teams had slightly higher t values in replications (ρ = 0.21, p = 0.05), as did groups in higher-ranked universities (replications (ρ = 0.18, p = 0.13), which also found lower effect size decreases between the original study and replication (ρ =-0.22, p = 0.07). Conversely, replicating teams with larger numbers of published articles found greater effect size decreases (ρ = 0.28, p = 0.008 for protocol-developing teams, ρ = 0.15, p = 0.15 for data collection teams). More experienced data collection teams were also more likely to consider the replication successful in their subjective assessments (ρ = 0.31, p = 0.003 for years since first publication and ρ = 0.24, p = 0.02 for number of publications). Once more, the multiplicity of tested predictors and non-independence between replications with overlapping teams make these correlations tentative at best.

### Self-assessment by labs and coordinating team

Protocol deviations occurred frequently in our sample and were registered by the coordinating team in 95% of individual replications (88% for MTT, 98% for PCR and 100% for EPM). On a scale of 1 to 5 (with 1 being no deviation and 5 being a deviation that invalidates the experiment as a direct replication), replications in the overall sample had a mean (± SD) rating of 1.9 ± 1 when evaluated by the lab and 2.5 ± 1.1 when rated by the validation committee (excluding replications where issues were found after initial validation, in which scores were not updated). Among experiments included in the primary analysis, these ratings were 1.7 ± 0.8 and 2.1 ± 0.8, respectively. When analyzing all experiments in an exploratory manner (see **Figure S8**), deviation scores by the committee were higher for animal (ρ = 0.42, p = p = 7.2 x 10^-7^) and EPM experiments (ρ = 0.4, p = 1.7 x 10^-6^) and lower for cell line and MTT ones (ρ =-0.49 p = 2.9 x 10^-9^). Protocol deviations also correlated with higher original coefficients of variation (ρ = 0.42, p = 8.1 x 10^-7^) and lower original standardized effect sizes (ρ =-0.36, p = 2.4 x 10^-5^) – probably due to confounding of these features with EPM experiments –, as well as with earlier publication years (ρ =-0.37, p = 1.4 x 10^-5^) and degree of challenge as assessed in the prediction survey (ρ = 0.31, p = 3.5 x 10^-4^).

Reasons for protocol deviations provided by labs are presented in **Figure 5**, with illustrative examples presented in **Table S22**. Most deviations were due to issues intrinsic to the experiment, such as the experimental model behaving differently than expected. Reasons related to the infrastructure of the lab and animal facility were also frequent, as were logistical problems involving suppliers or regulatory requirements. A smaller fraction of deviations was due to deliberate choices or errors by the lab performing the replication. An assessment of general difficulties by participating researchers (**Figure S9** and **Table S23**) placed COVID-19 restrictions, delayed delivery of reagents, and difficulties with the experimental model as the top challenges faced by labs.

**Figure 5.**
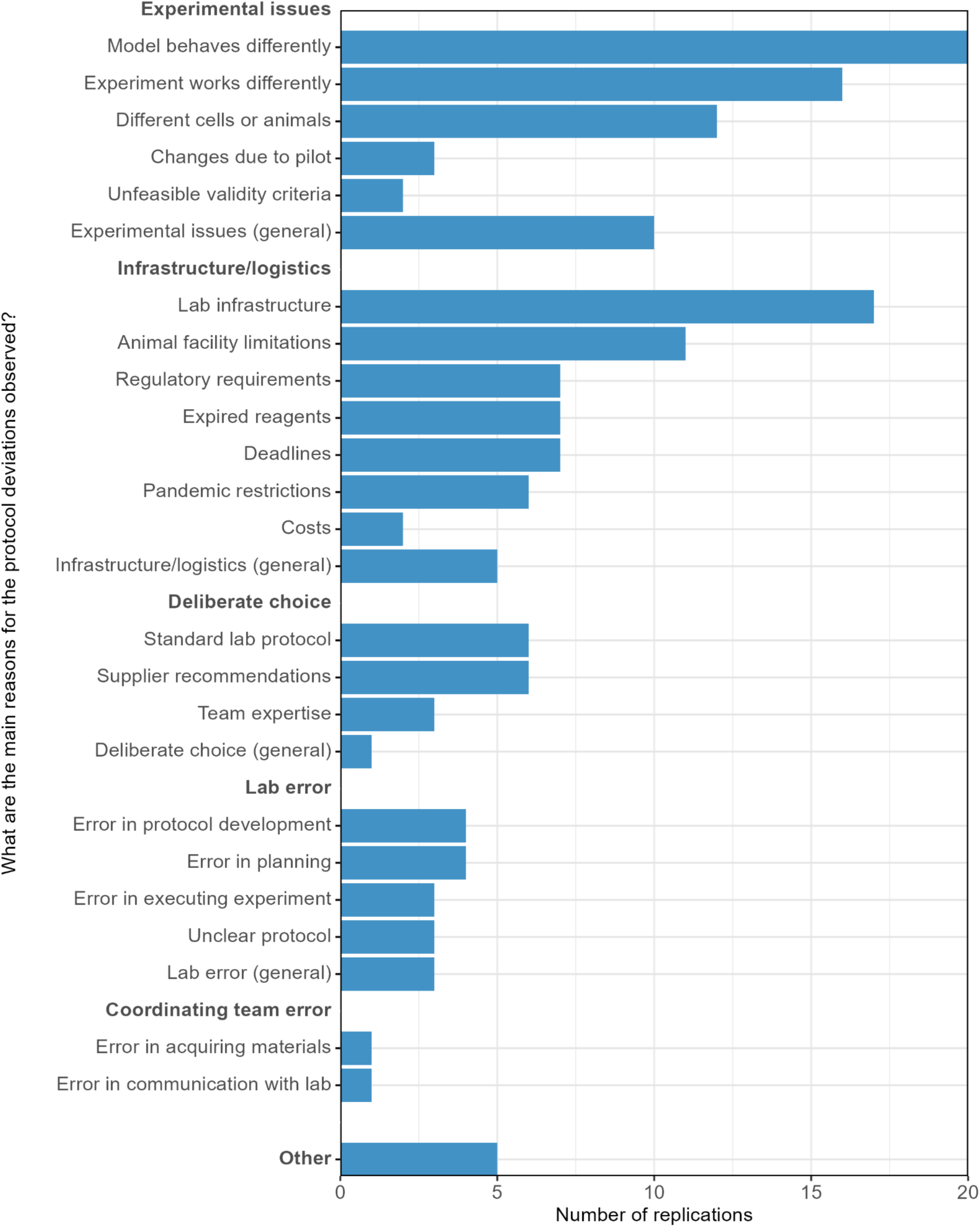
Reasons for protocol deviations. Reasons were provided as open answers by the lab and categorized by the coordinating team as described at https://osf.io/5gjb7. They are divided into general categories (bold) and subcategories. Examples for each category are shown in **Table S22**.

The coordinating team also assessed the obstacles faced by the project, with top-ranked difficulties shown in **Table S24** and a complete list available at https://osf.io/q76vj. Lack of common terminologies for describing aspects of experiment design, problems with reagent delivery, poor methodological descriptions in the original articles, and experimental models not working as expected were considered the most relevant difficulties. These were grouped into five categories encompassing the key challenges faced by the project (**Table 4**). These challenges, along with associated opportunities for improvement, will be discussed more extensively in future publications.

**Table 4.** Challenges in performing a large-scale multicenter replication of experiments from Brazilian biomedical science. Main categories of challenges faced by the project, based on the coordinating team’s assessment of an initial list of difficulties (see **Table S24** and https://osf.io/q76vj).

| Category | Examples |
| --- | --- |
| Lack of standardized terminologies for experimental design and reporting | Unstandardized/incomplete descriptions of methods and results in original articles.<br>Discrepancies in terminology used to describe issues such as experimental units and controls by replicating labs. |
| Tension between replicating protocol exactly and getting methods to work | Lack of consensus between essential and adaptable steps in protocols.<br>Experimental models behaving differently across labs.<br>Difficulties in preregistration in the absence of pilot experiments.<br>Difficulties in establishing criteria for methodological validity. |
| Multicenter project management | Communication difficulties between the coordinating team and participant labs<br>Lack of formal training in project management by the coordinating team.<br>Participant labs not used to external supervision or standardization of procedures.<br>Difficulties in asynchronous review of protocols and data.<br>Lack of data management standards among participant labs. |
| Logistics and infrastructure | Difficulties in obtaining licenses with regulatory agencies.<br>Inefficiency of national suppliers in delivering reagents.<br>Limitations in animal facilities and general laboratory infrastructure.<br>High prevalence of junior scientists in participating teams. |
| General difficulties for large-scale projects | Difficulties in anticipating loss of labs and experiments during the project.<br>Difficulties in data management and statistical analysis of complex data.<br>Difficulties in maintaining documentation practices throughout the project. |

## Discussion

In this large multicenter replication of published biomedical experiments from Brazil, replication rates in our primary analysis varied from 20 to 44%, depending on the criteria used to define replicability. These numbers are in a similar range as previous prospective estimates from the biomedical literature (Begley & Ellis, 2012; Errington, Mathur, et al., 2021; Prinz et al., 2011; Scott et al., 2008; Steward et al., 2012). Among these, our study is unique in trying to assess a representative sample of articles by randomly selecting experiments from a wide range of publications based on methods, rather than filtering by research area, importance or citation counts. It is also the first attempt to evaluate replicability within a specific country, in order to provide data that can be of more direct use for local funders, institutions and researchers. That said, the absence of similar studies elsewhere precludes comparisons with other regions.

When compared to originally positive experiments in the Reproducibility Project: Cancer Biology (RP:CB) (Errington, Mathur, et al., 2021) – perhaps the closest counterpart to our study – replication rates in our primary analysis were lower when measured by statistical significance in the same direction (20% *vs*. 39%) or inclusion of the replication effect size in the original 95% confidence interval (29% *vs*. 43%). Correlations between effect sizes were also lower in our study (Spearman’s ρ = 0.35 *vs*. 0.47), while effect size decreases were similar in magnitude (85% in standardized effect sizes in RP:CB, 90% in relative differences in our study). Multiple factors could account for the differences, including sampling strategy (general literature *vs*. highly cited articles), experimental selection (first significant experiment *vs.* most relevant ones), replication facilities (academic labs *vs*. commercial services), protocol development (with *vs*. without the participation of original authors), subfield (general *vs*. cancer biology) and region of origin (Brazil *vs*. worldwide).

Our study is also unique in employing a multicenter replication strategy, allowing us to assess the extent to which interlaboratory variability may account for discrepancies between the original study and its replications. Differences in effect sizes between the original study and replication were larger than those among replications, suggesting that irreplicability is partly due to factors that are intrinsic to published articles, such as publication bias, selective reporting, low statistical power and incentives for positive results (Smaldino & McElreath, 2016). That said, differences between replications were also large in many cases. As replicating labs were blind to the original result and had no obvious incentives towards particular results, heterogeneity between replications is more likely due to differences in experimental conditions or in how experiments were conducted. These may include protocol adaptations, different levels of adherence to the original protocol, or undetected errors by replicating labs. The finding of significant interlaboratory variability is also consistent with previous attempts to replicate identical protocols across multiple sites for animal (Crabbe et al., 1999) and cell line studies (Elliott et al., 2017; Niepel et al., 2019).

Within the biomedical literature, our study is also one of the few to assess predictors of replicability. Although the power of this analysis was limited by the small number of successful replications, MTT and cell-line experiments exhibited smaller decreases in effect sizes between the original and replication than EPM and animal experiments. Reasons for irreplicability also seemed to differ between methods. In PCR experiments, non-significant results in some experiments were likely explained by coefficients of variation being higher in replications than in original studies, leading to low statistical power. For the other methods, power was usually adequate and non-significant replication effects were closer to zero, suggesting that original results are more likely to represent false-positive or ungeneralizable results.

Although researchers’ predictions overestimated replicability, they had reasonable success in forecasting the evidence strength of replications as measured by t values; still, prediction accuracy was lower than that reported by previous replication projects in psychology and the social sciences (Gordon et al., 2021). This finding might be expected, given that the replicability of findings in biomedical science is probably less intuitive than that of psychology studies, for which prediction accuracy is above chance even among laypeople (Hoogeveen et al., 2020). Assessments of how challenging the replications would be, meanwhile, correlated both with experimental complexity measured by text-based assessment of the protocols (Fanelli et al., 2025) and with greater effect size decrease between the original result and replication. Researcher justifications for predictions and analyses of whether prediction markets can improve accuracy will be explored in further work.

Other features of the original article were generally uncorrelated with replication outcome, although large rates of publications by the last author were associated with lower replicability, suggesting that incentivizing publication volume may be counterproductive for the reliability of results. Journals and university rankings were not predictive of replicability, in contrast with a recent retrospective analysis of *Drosophila* immunity studies (Lemaitre et al., 2026), although the focus on randomly selected Brazilian articles led our sample to mostly include low impact factor journals. Counterintuitively, greater use of measures to control risk of bias in animal experiments, such as randomization and blinding, was correlated with greater effect size decrease in the replication. Although this is likely to be a chance finding, as both sample size for the analysis and plausibility are low, it suggests that replicability in lab biology – as well as predictions about it – may rely more on the complexity of experiments than on traditionally used proxies for methodological rigor.

Our analyses have multiple caveats, most of which are due to difficulties faced by participating labs performing replications. Twenty-one percent of planned replications ended up not being performed, either because materials could not be acquired (5%) or because the labs to which they had been assigned left the project or could not meet its deadlines (15%). This is a much smaller fraction than the 74% unfinished replications in RP:CB (Errington, Denis, et al., 2021), likely due to our selection of simpler experiments. On the other hand, our validation process had a large impact on our sample, excluding 37% of replications from the primary analysis. Interestingly, this filtering process did not have a large impact on replication rates, as included and excluded replications had similar rates of success. This may be explained by the fact that, while some issues raised during the validation process would be expected to lower replicability (e.g. deviations from the original protocol), others could artificially raise it (e.g. insufficient biological variation among experimental units or biases in randomization).

Loss of individual replications also limited statistical power for many experiments, even though we aimed for high power in sample size calculations (95% power to detect the original effect in each individual replication) to account for the likelihood of effect size inflation (Ioannidis, 2008; van Zwet et al., 2023). Power was further constrained by the fact that coefficients of variation were higher in replications than in original studies, with a 4-fold median difference for PCR experiments. This may either indicate that replicating labs had more difficulty in controlling technical error or that variability was underestimated in the published results, due to elimination of outliers (Gress et al., 2018; Holman et al., 2016), incorrect labeling of error bars (Cumming et al., 2007; Vaux, 2004), or insufficient biological variation among experimental units (Lazic et al., 2018; Vaux et al., 2012). Filtering the sample by achieved power (considering the original relative effect and the observed variability) suggests that higher variability in replications biased replication rates based on statistical significance downwards; nevertheless, even when considering only experiments with adequate power, replication rates remained between 26 and 38%.

For experiments with high agreement among replications, it is reasonable to assume that the reasons for irreplicability are more likely to lie in the published results. Nevertheless, as we replicated individual comparisons between groups, lack of replication success does not apply to the whole article, or even to the whole experiment from which the comparison was drawn. Due to our choice of selecting the first comparison with a significant difference in an article, it was not infrequent to select an initial experiment or a low treatment dose, meaning that an unsuccessful replication does not necessarily call the main conclusions of a study into question.

More importantly still, low agreement among replications and/or a lower number of replications than originally planned limited our ability to judge many of the original experiments. Our results should thus be thought of as naturalistic estimates of real-life academic labs trying to replicate individual experiments from the published literature. That said, due to our crowdsourced, country-based design, the population of labs that performed replications was largely similar to the one that produced the original results. Thus, whether the original estimate or the replication one is a more accurate description of the phenomenon under study does not change the conclusion that Brazilian researchers have difficulties replicating findings from other research groups within the same community.

In fact, the observation that most labs faced difficulties in following preregistered protocols and controlling experimental variability provided an opportunity to glimpse at possible causes for replication failures. While all we can say about the original results is how closely they were replicated by our labs – which provides little information about the reasons for failed replications – our data collection and review process allowed an extensive audit of how closely replication protocols were followed by participant labs. This was complemented by a self-assessment process that allowed us to investigate relevant difficulties and opportunities for improvement.

Deviations from preregistered protocols were present in most replications. They occurred most commonly due to issues that were intrinsic to the experiment – such as difficulties with growing cells in the conditions reported in the original publication, or with performing experimental procedures as stated in the protocol. In many cases, labs reported that these issues could not have been anticipated without direct experience with the experimental model. Similar limitations were observed in the RP:CB, in which two thirds of preregistered protocols required modifications after experimental work had begun (Errington, Denis, et al., 2021).

This should be considered when interpreting replication rates, as difficulties in adequately implementing a method or experimental model are likely to decrease the probability of replication success. Thus, “one-shot” replication projects such as our own, in which extensive efforts are made to define the protocol in advance, are different from “best-shot” attempts at replication that allow for tinkering with the protocol over the course of multiple replication attempts – which are arguably closer to how experiments are typically replicated by academic labs (Guttinger, 2019). While the former are likely to underestimate replicability due to the difficulty in defining methods upfront, the latter can overestimate it due to the active effort in obtaining results that are close to the original.

The challenge of prespecifying methods also poses limits to the applicability of protocol preregistration in laboratory biology – or in any other area in which data collection depends on complex methods. Although we believe that preregistration of experimental methods is worthwhile, it may be more useful when performed after pilot experiments have been carried out, in order to reduce the need for deviations. Preregistering analysis methods poses fewer difficulties when data is relatively simple, as in our experiments; nevertheless, more data-intensive methods can involve steps of data filtering or complex analytical decisions (e.g. Carp, 2012) that may be equally hard to specify in advance.

Our attempts to establish predefined inclusion/exclusion criteria to assess the validity of individual samples or experiments (Neves & Amaral, 2020) were also less successful than we had expected. Labs had difficulties in establishing these criteria in advance, frequently setting very high standards for markers, such as RNA purity, that turned out to be difficult to achieve and led them to break their own criteria. An independent validation process performed after the experiment turned out to be more feasible for assessing methodological validity and may be more compatible with the reality of academic labs. Importantly, this process was blind to the replication results, in order to prevent replication outcomes from contaminating judgments about methodological adequacy. Nevertheless, some methodological issues went undetected by this process and were identified only when results were actively discussed with participating labs.

Other reasons for protocol deviations included infrastructural problems, such as difficulties with the delivery of materials, cells or animals on time, limitations in access to equipment, and lack of personnel to perform replications. The COVID-19 pandemic proved to be a particularly complex challenge, as many labs and institutions were closed for extended periods, and even after normal activity was resumed, changes in replicating teams and lab priorities meant that many experiments ended up being performed in non-ideal conditions. Unsurprisingly, this was the most frequently reported difficulty by participants in our project evaluation survey.

All that said, there remains a fraction of protocol deviations that had no particular reason, in which labs simply made a different methodological choice from the preregistered one. This suggests that the idea of preregistration as a commitment to a particular methodology is still poorly understood by biomedical researchers. Most of them seem to consider protocols as general guidelines, assuming – perhaps correctly – that many steps will change after the experiment begins, and sometimes preferring vague descriptions over precise specifications to allow flexibility in the process. In some cases, this also led to a lack of clarity when developing protocols, leading to misunderstandings when experiments were performed by a different researcher.

These communication issues were particularly evident when determining the appropriate experimental unit in cell culture experiments. Although this was defined by the coordinating team in the protocols as different passages of a cultured cell line, or as primary cultures from independent pools of animals, many labs opted for other ways to define biological replicates. Not all adaptations were accepted, leading many replications to be excluded from our primary analysis. Moreover, merely understanding what experimental unit was used by a particular lab could be challenging, due to the lack of standardized terminology to define this among researchers (Lazic et al., 2018; Vaux et al., 2012). To a lesser extent, this also applied to other aspects of experimental design, such as positive and negative controls, blanks and randomization, which were described using distinct terminology by individual labs. This suggests that the adoption of consensus taxonomies that systematically and unambiguously describe these concepts could potentially improve reproducibility.

A final lesson learned from this project is that organizing large-scale initiatives in laboratory biology is hampered by the lack of experience of academic labs with working in coordinated fashion. While collaboration in biomedical research is frequent, the culture of repeating the same experiment across labs and standardizing procedures is confined to specific research fields (Brancato et al., 2024) or projects (Lyden et al., 2022). As a striking example of this limitation, a recent systematic review of the biomedical literature found only sixteen examples of multicenter animal studies (Hunniford et al., 2023). Consequently, data collection and management standards by labs are idiosyncratic, and harmonizing, reviewing and analyzing data from different sources can be challenging. Moreover, managing large-scale projects is a skill for which researchers receive little training, and being externally managed or supervised is also an unusual experience that can be met with resistance (Maysami et al., 2016). If large-scale, crowdsourced research is to flourish in lab biology, cultural changes are important to capacitate the academic workforce for such collaborative endeavors (Amaral & Neves, 2021; Coles et al., 2022; Uhlmann et al., 2019). Possible solutions in this direction will be elaborated in a future article dedicated to these issues.

As a final reflection, our project leaves contributions beyond the results reported here. Our quantitative data describing replications are fully shared and open to scrutiny. Similarly, most of our textual data – including protocols, final notes, justifications for researcher predictions and other documents – are also open and remain to be analyzed in more depth. The collaborative nature of the project has constituted an important learning experience for the many researchers involved, including many early-career scholars, and the visibility we received (e.g. de Oliveira Andrade, 2019, 2025) has cast a spotlight on replicability and reproducibility in academic debates across the country. Finally, we leave spinoffs such as the Brazilian Reproducibility Initiative in Preclinical Systematic Review and Meta-Analysis (BRISA) (http://en.reprodutibilidade.bio.br/brisa), an ongoing collaboration to train biomedical researchers in systematic review methods, and the Brazilian Reproducibility Network (http://en.reprodutibilidade.org), a multidisciplinary organization to promote reproducible research practices across the country. We expect this legacy to help place reproducibility at the forefront of Brazil’s national research agenda.

## Notes

https://osf.io/6av7k/

https://doi.org/10.7910/DVN/ZJRDIV

https://github.com/BrazilianReproducibilityInitiative/bri-analysis

### Competing Interest Statement

Kleber Neves and Clarissa Carneiro were hired by the study's funder (the Serrapilheira Institute, a philantropic organization with no financial stake at the results) midway through the project and were thus employed by it during data analysis and manuscript preparation.

### Summary of Updates

- Data corrections were performed in 26 replications and validation decisions were reviewed for 7 replications after systematic checking of data with labs via online meetings (see list of data changes at https://osf.io/4tu2v). - The main analysis for MTT experiments was changed to include pairing between experimental units in the treated and control group as the default, as absolute optical density values were found not to be commensurable across units. The original approach is maintained as a sensitivity analysis. - Pairing is now implemented by normalization to a particular experimental unit rather than in the meta-analysis methods alone. - A sensitivity analysis using the mean of all experimental units (normalized by controls in each replication) rather than meta-analysis was added. - Various changes were made throughout the text and figures in response to reviewer comments at a journal. A list of these (as well as the responses to comments) can be found at https://osf.io/6av7k/r5y9x

### Funding

- Instituto Serrapilheira
- Fundação Carlos Chagas Filho de Amparo à Pesquisa do Estado do Rio de Janeiro: E-26/200.824/2021
- Conselho Nacional de Desenvolvimento Científico e Tecnológico (CNPq): 310813/2021-2

