## Supplementary Figures and Tables for "Estimating the replicability of Brazilian biomedical science"

**Estimating the replicability of Brazilian biomedical science - Supplementary Material**

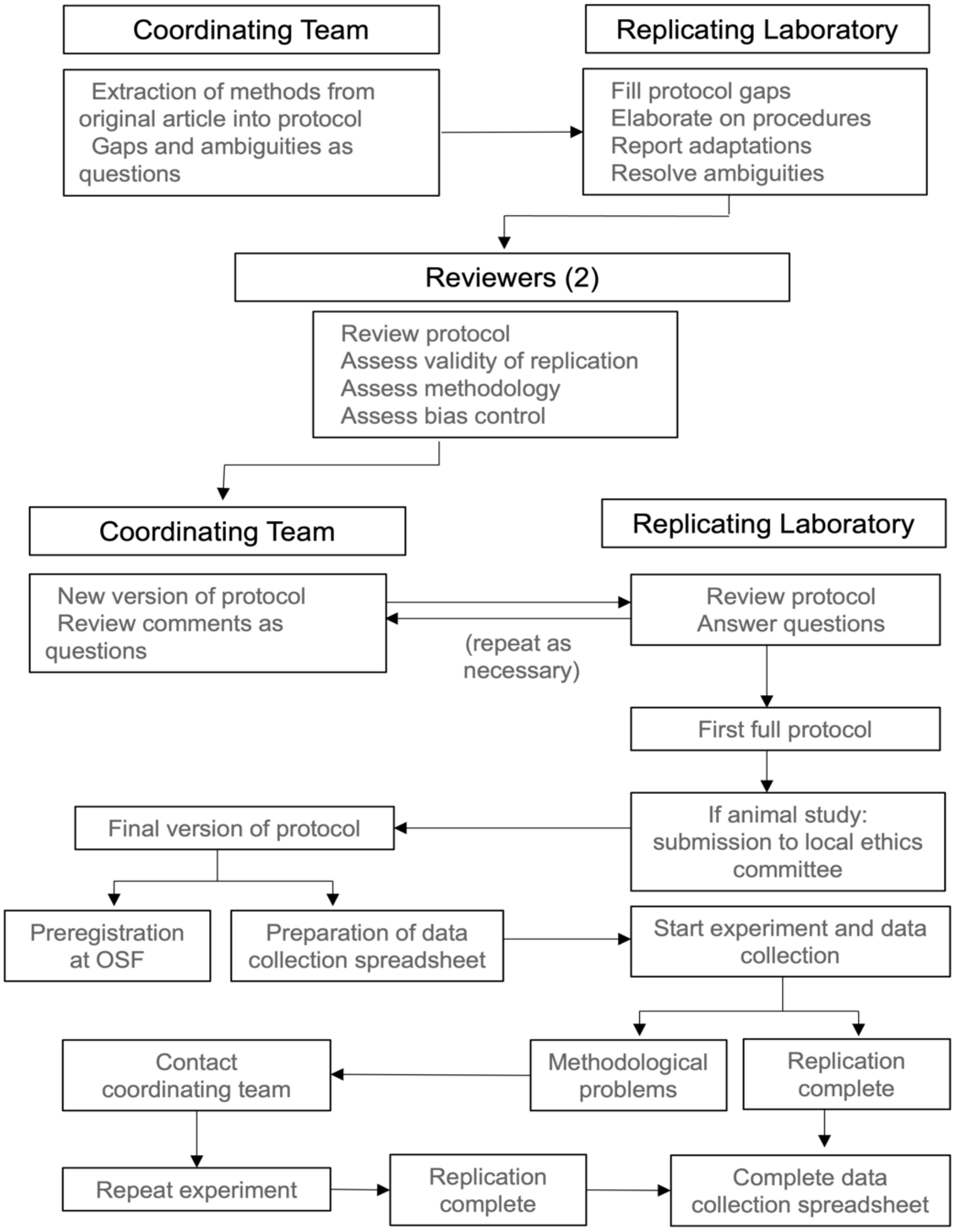


**Figure S1. Summary of protocol development and replication**. Methods from the original article were extracted by the coordinating team and sent to the replicating lab, with gaps and ambiguities formulated as questions. After these were answered, protocols were reviewed by two independent reviewers, with comments also sent to the lab as questions. When these were resolved, the protocol was registered (after evaluation by an ethics committee in the case of animal experiments). Data collection spreadsheets based on the protocol were developed and sent to the lab, which was charged with performing the replication, contacting the coordinating team when needed if methodological problems arose.

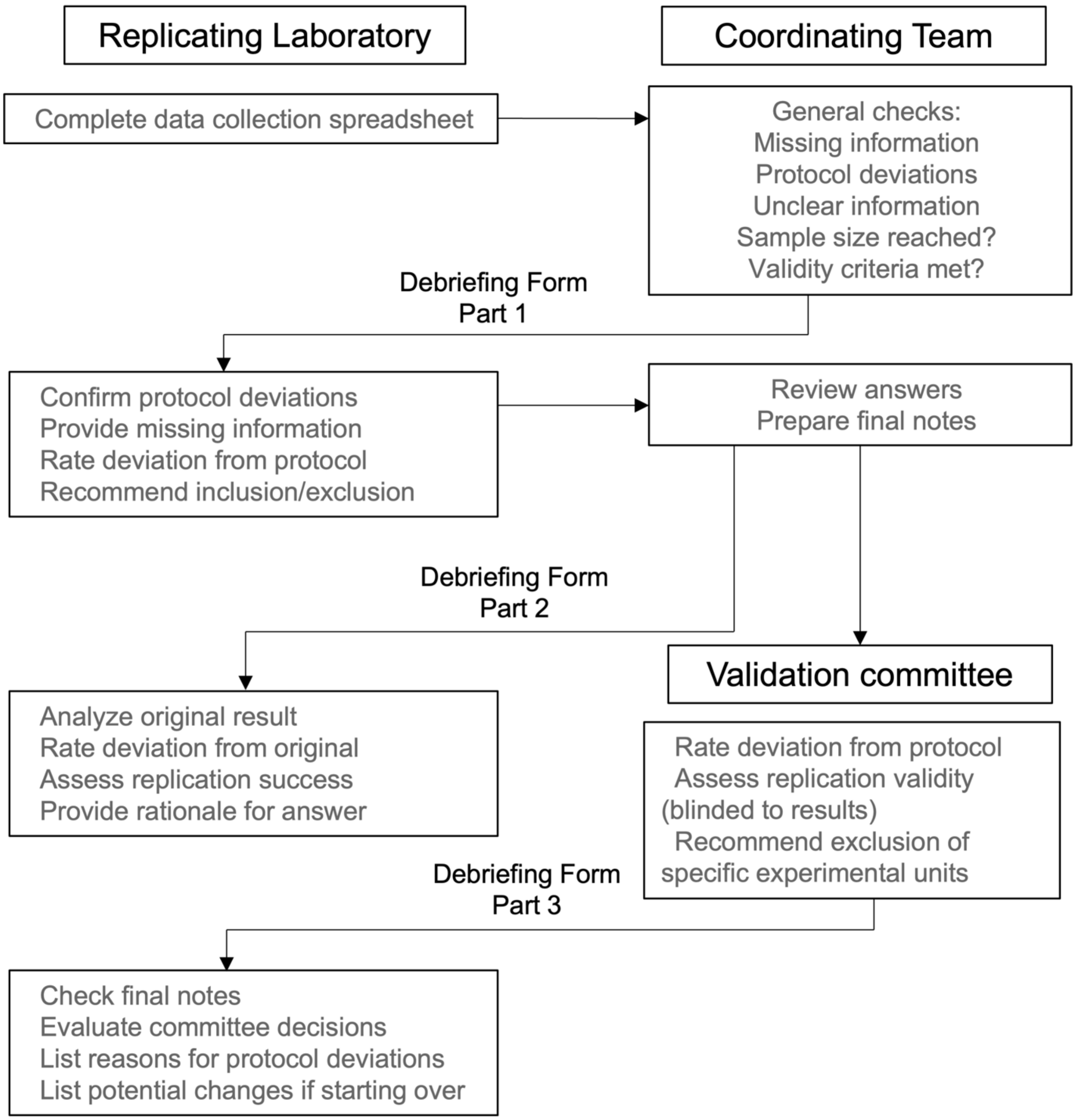


**Figure S2. Summary of experimental debriefing and validation**. After data collection spreadsheets were complete, they were reviewed by the coordinating team, which checked protocol adherence and sent questions and requests for missing information to the replicating labs. Based on their answers, final notes documents were developed and sent to the validation committee, which evaluated protocol deviations and decided whether the experiment should be included in the primary analysis. Meanwhile, replicating labs received the original article and were asked to provide their assessment of replication success. Finally, they received the validation committee's assessment and were asked about their agreement with it, as well as about reasons for protocol deviations and ways in which they could have been prevented. For validation decisions that were later reviewed by the committee, questions about agreement were resent to the lab and only the final answer was considered.

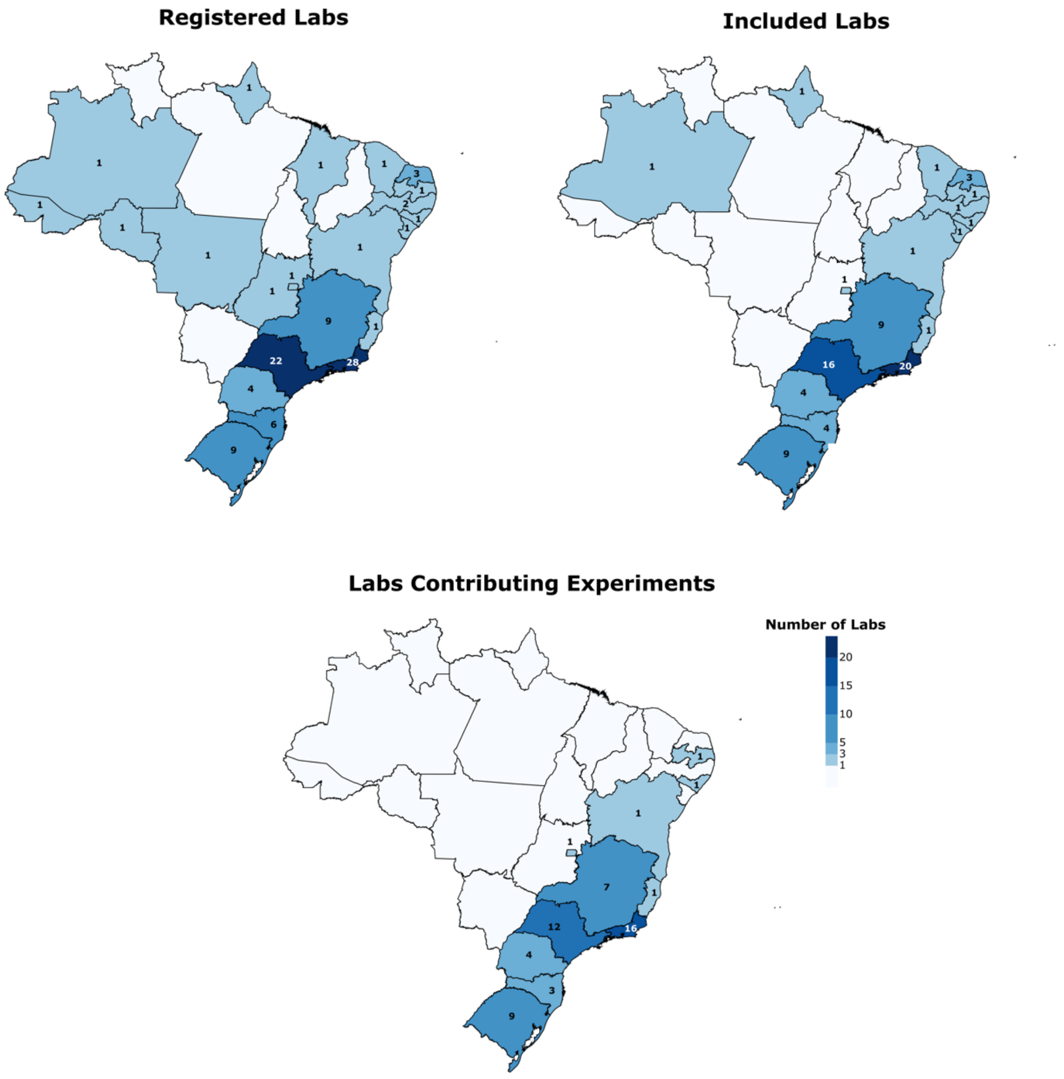


**Figure S3. Geographic distribution of registered and participating labs.** Maps show the number of labs per Brazilian state registering for the project (top left), receiving experiments for replication (top right) and contributing data (bottom). These data are presented in tabular form in **Table S1**.

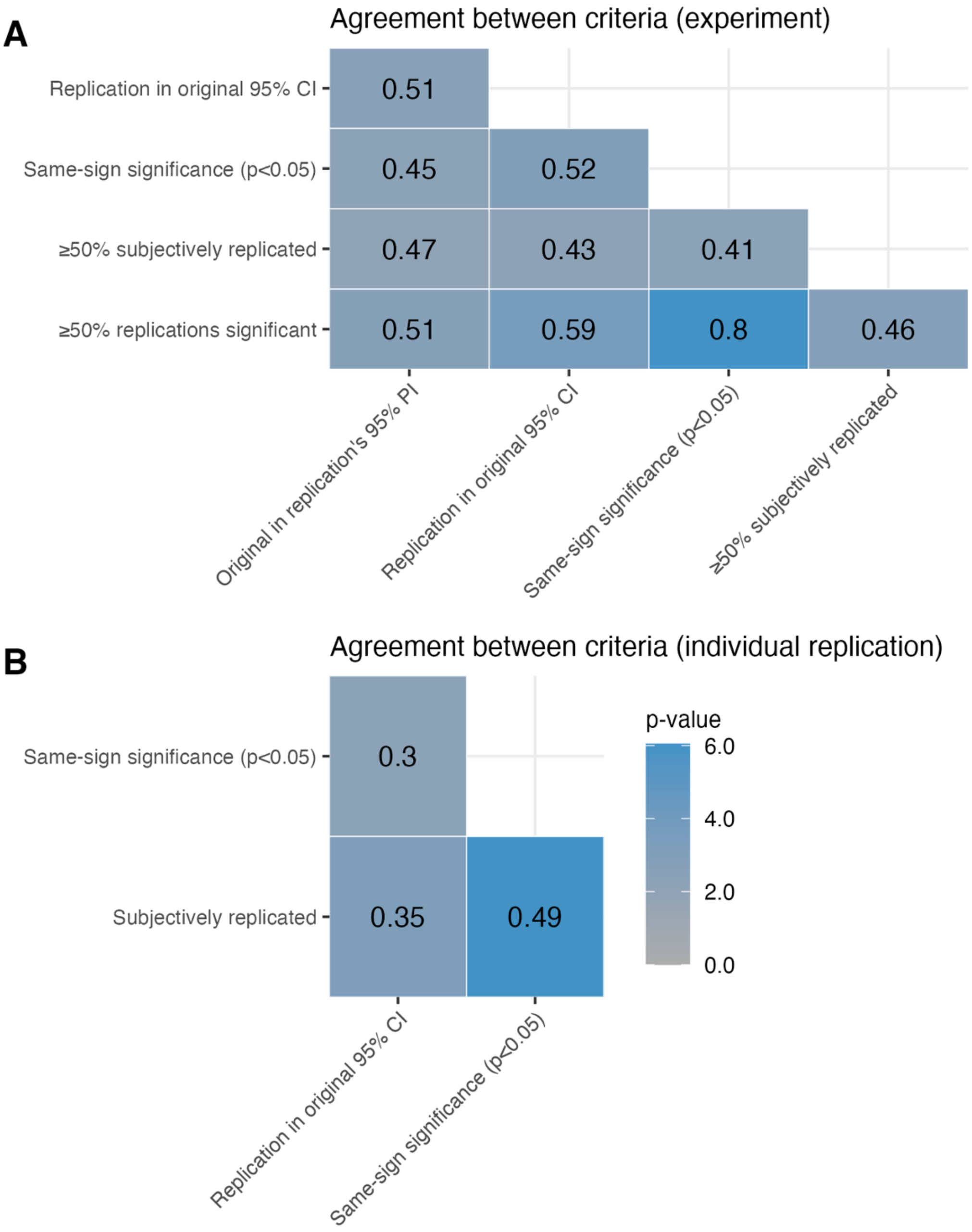


**Figure S4. Agreement between replication criteria.** Matrices show Cohen's kappa coefficients for each pair of replication criteria at the level of experiments (**A**) and individual replications (**B**), with the color scale indicating *p* values for agreement. Fleiss' kappa coefficients are 0.54 ($p = 8.3 \times 10^{-27}$) for the 5 experiment-level criteria and 0.38 ($p = 5.8 \times 10^{-10}$) for the 3 replication-level criteria. Criteria are the same as in **Table 1**, with same-sign significance for experiments based on a fixed meta-analysis estimate, effect size comparisons based on random-effects meta-analysis, and analysis using the t distribution based on the number of experimental units. Agreement with the 95% prediction interval criteria and Fleiss kappa calculations only take into account experiments with more than one replication. PI, prediction interval, CI, confidence interval.

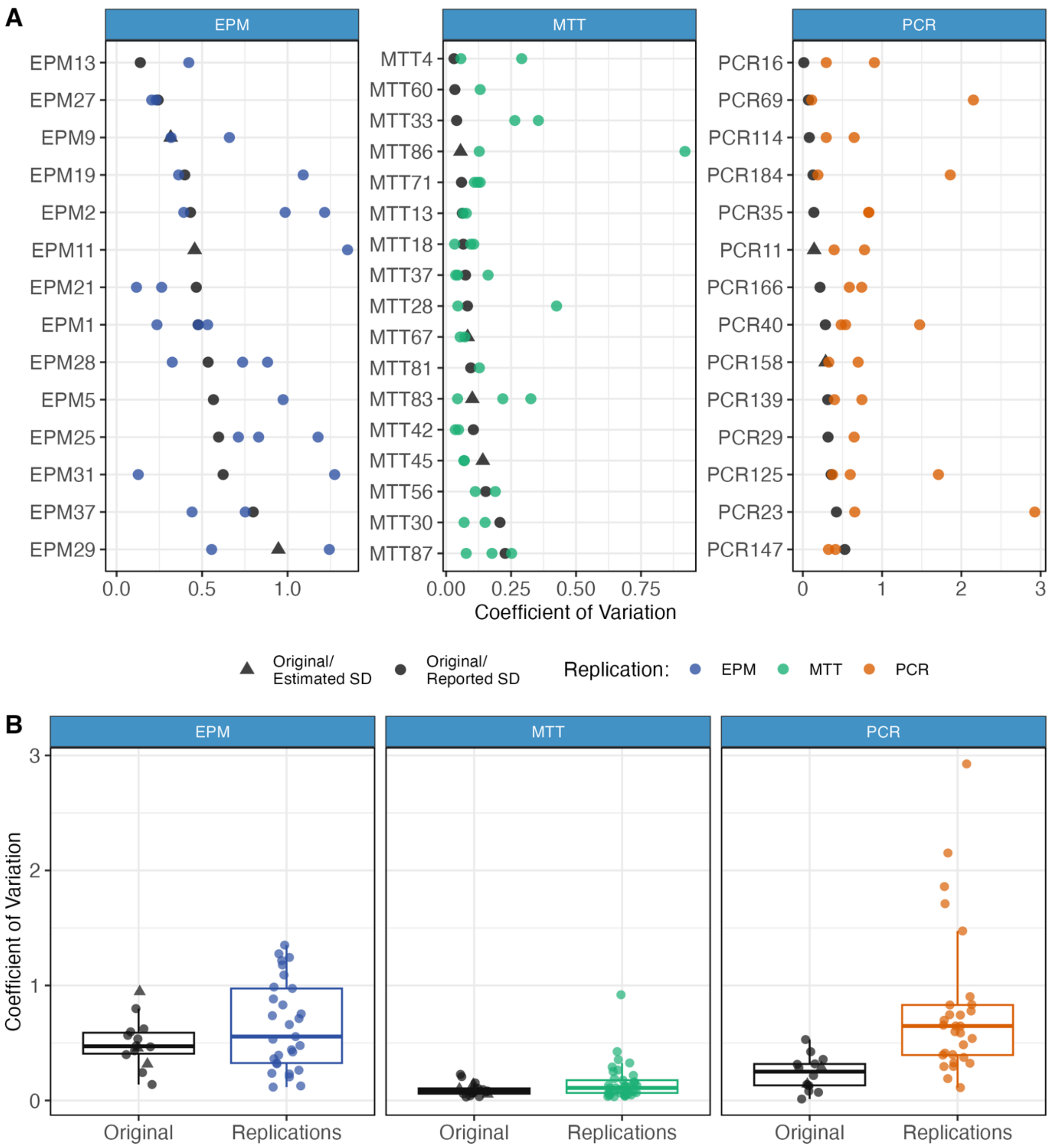


**Figure S5. Coefficients of variation in original experiments and replications. (A)** Coefficients of variation (CVs) in original experiments (dark) and individual replications (light) for EPM (blue), MTT (green) and PCR (orange), ordered by the original CV. X-axes represent CVs, calculated as the pooled standard deviation divided by the mean of both groups. Triangles indicate experiments that did not report exact SDs, in which they were estimated as the mean of the sample size range provided. PCR experiments use coefficients of variation between linearized expression values (usually $2^{-\Delta\Delta Ct}$ for real-time PCR and relative expression for conventional PCR). Paired experiments use CVs based on SDs after paired normalization. **(B)** Group comparison between original and replication results for each method. Results were compared with Wilcoxon's signed rank test, yielding $p = 0.05$ for EPM, $p = 0.15$ for MTT, $p = 2.4 \times 10^{-4}$ for PCR and $p = 1.3 \times 10^{-5}$ for all methods.

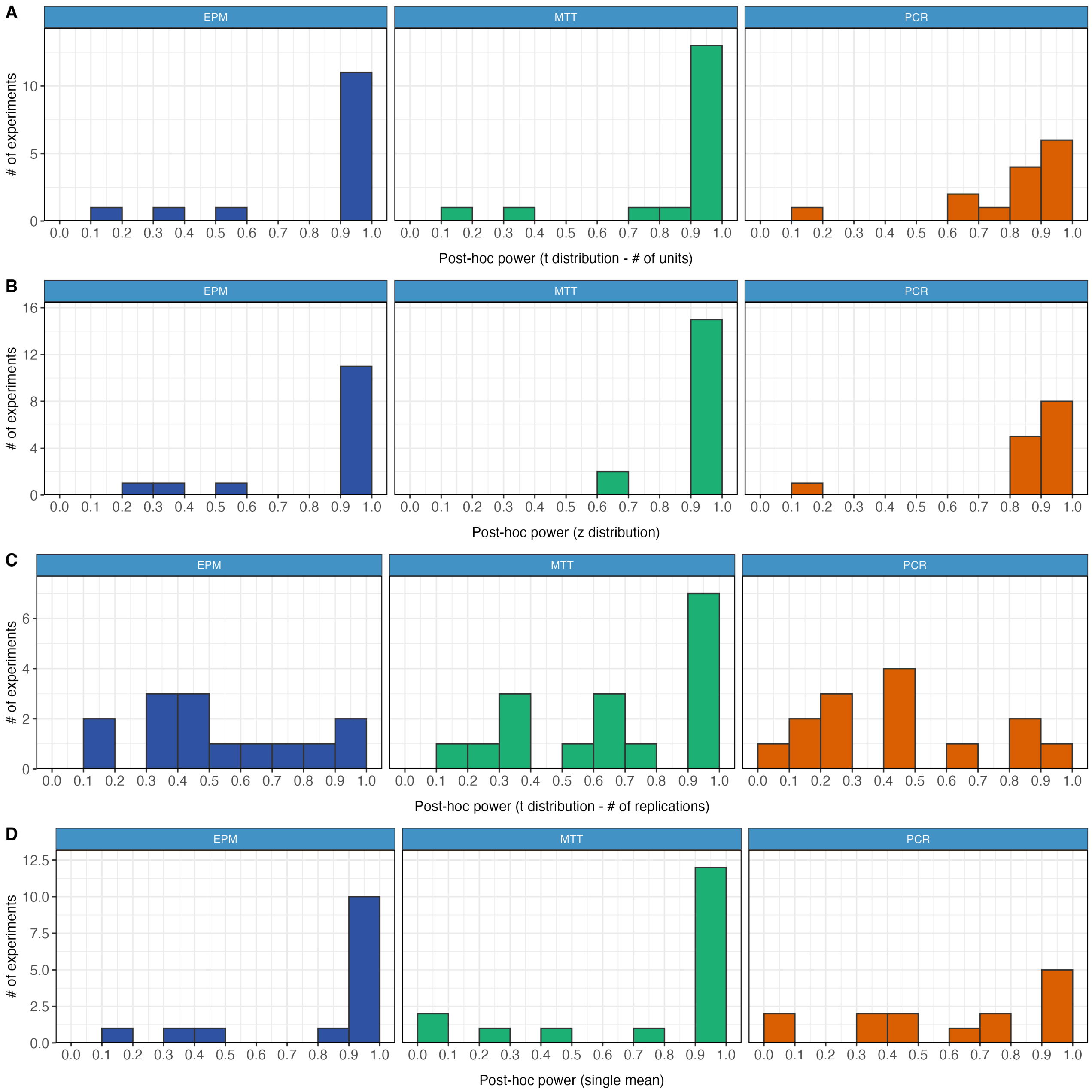


**Figure S6. Distribution of *a posteriori* power.** Histograms show the distribution of *a posteriori* power (calculated using the original relative difference and the variability achieved in replications) for EPM (blue), MTT (green) and PCR (orange) experiments included in the primary analysis using **(A)** the z distribution (first row), **(B)** the t distribution using the number of units (second row), **(C)** the t distribution using the number of replications (Knapp-Hartung approach, third row) and the single mean approach (fourth row). X-axes show the percentage of successfully replicated experiments using the same-sign significance criterion in a fixed-effects meta-analysis for 1,000 simulations of random data using the original ratio of means and the standard deviation observed in each replication after normalization by control values. Power is greater on average for EPM and MTT experiments than for PCR ones, with 10 experiments (3 EPM, 3 MTT, 4 PCR) failing the 80% threshold in the primary analysis. Power is greater with the z distribution, lower with the single mean approach, and lowest with the Knapp-Hartung method, which causes most experiments to be underpowered due to the low number of replications.

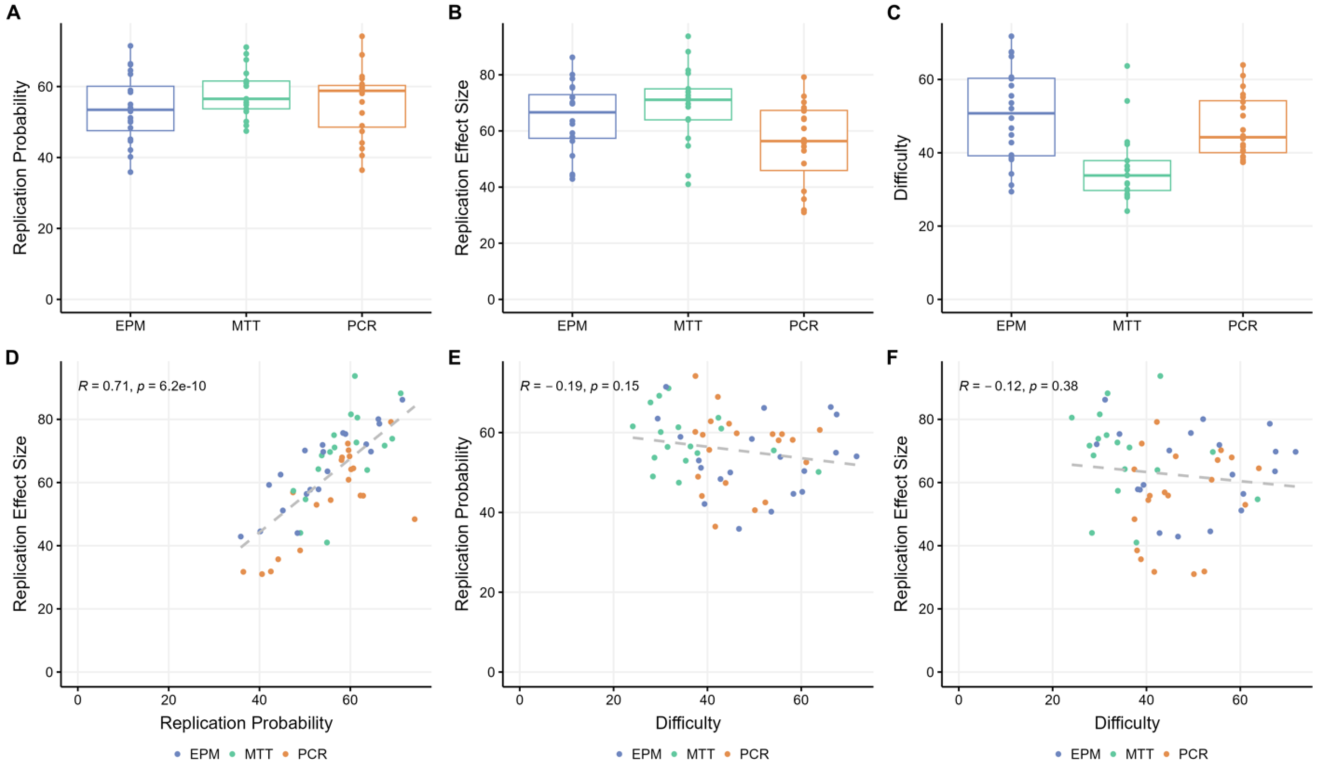


**Figure S7. Results of prediction surveys.** Box plots show the distribution of predictions for the 20 experiments using EPM (blue), MTT (green) and PCR (orange) concerning **(A)** probability of replication success (%), **(B)** predicted replication effect size (as % of the original relative effect) and **(C)** technical difficulty of the experiment. Each data point represents the mean of survey answers for that particular experiment by independent researchers (n=30 for EPM, 22 for MTT, 18 for PCR). Scatter plots show Pearson correlations between **(D)** replication probability and predicted effect size ($r = 0.71$, $p = 6.2 \times 10^{-10}$), **(E)** technical difficulty and replication probability ($r = -0.19$, $p = 0.15$) and **(F)** technical difficulty and predicted effect size ($r = -0.12$, $p = 0.38$).

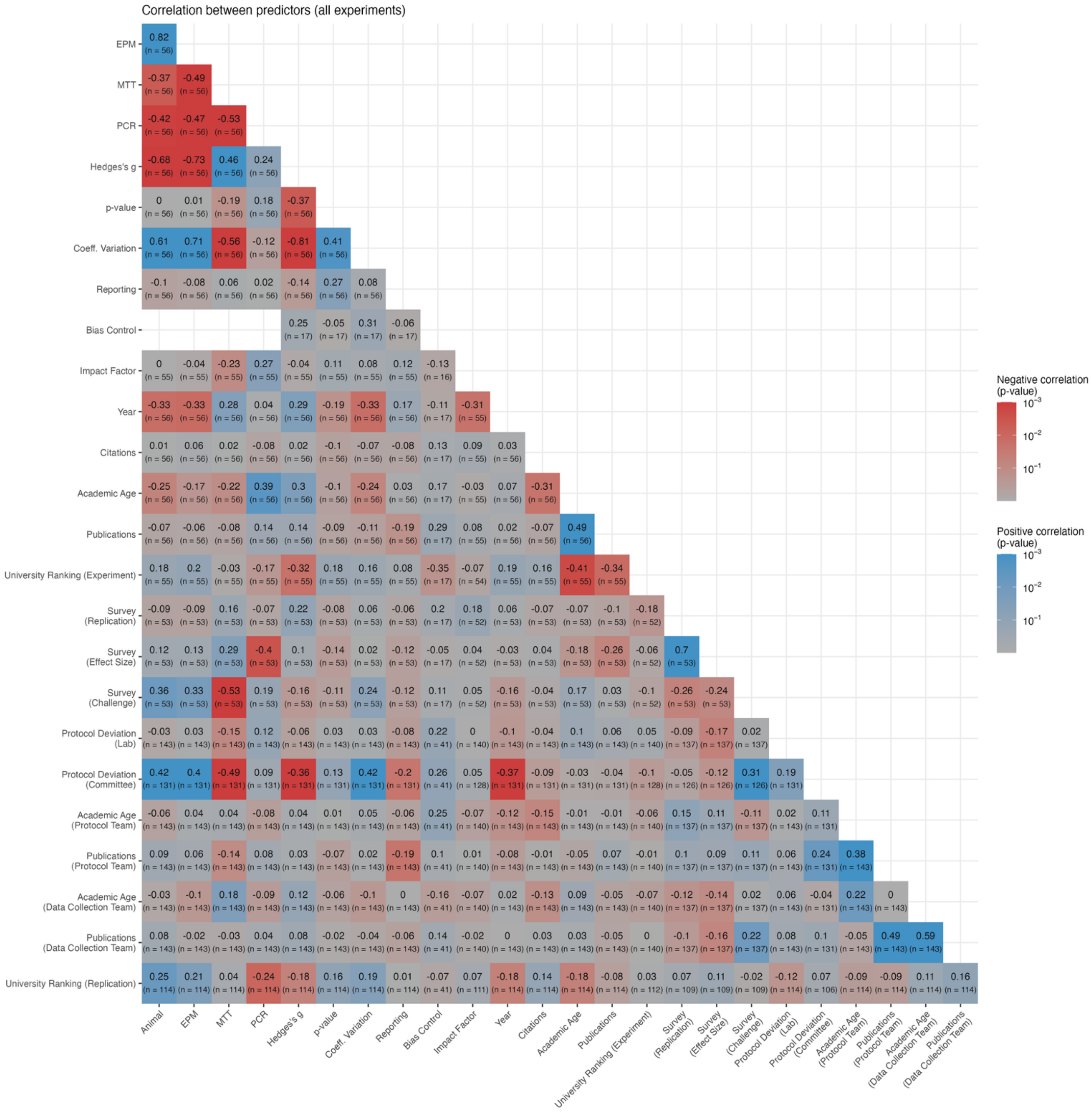


**Figure S8. Correlation between predictors of replicability.** Correlation matrices show values for Spearman's ρ between predictors of replicability at the level of both experiments and replications, with the color scale indicating *p* values. All experiments with data are used in the analysis. Predictors are the same as those shown in **Figure 4** and are described in detail at https://osf.io/9rnuj. Sample size is determined by the number of experiments for correlations between experiment-level predictors and by the number of replications for those that involve replication-level predictors. Experiments that were rediscussed with the validation committee after revision are not included in protocol deviation committee score analysis, as they were not rescored after review. Bias control measures are only used for EPM experiments in animals, and thus do not correlate with model or method. Sample size for correlations with university ranking is lower because some labs were not based in universities.

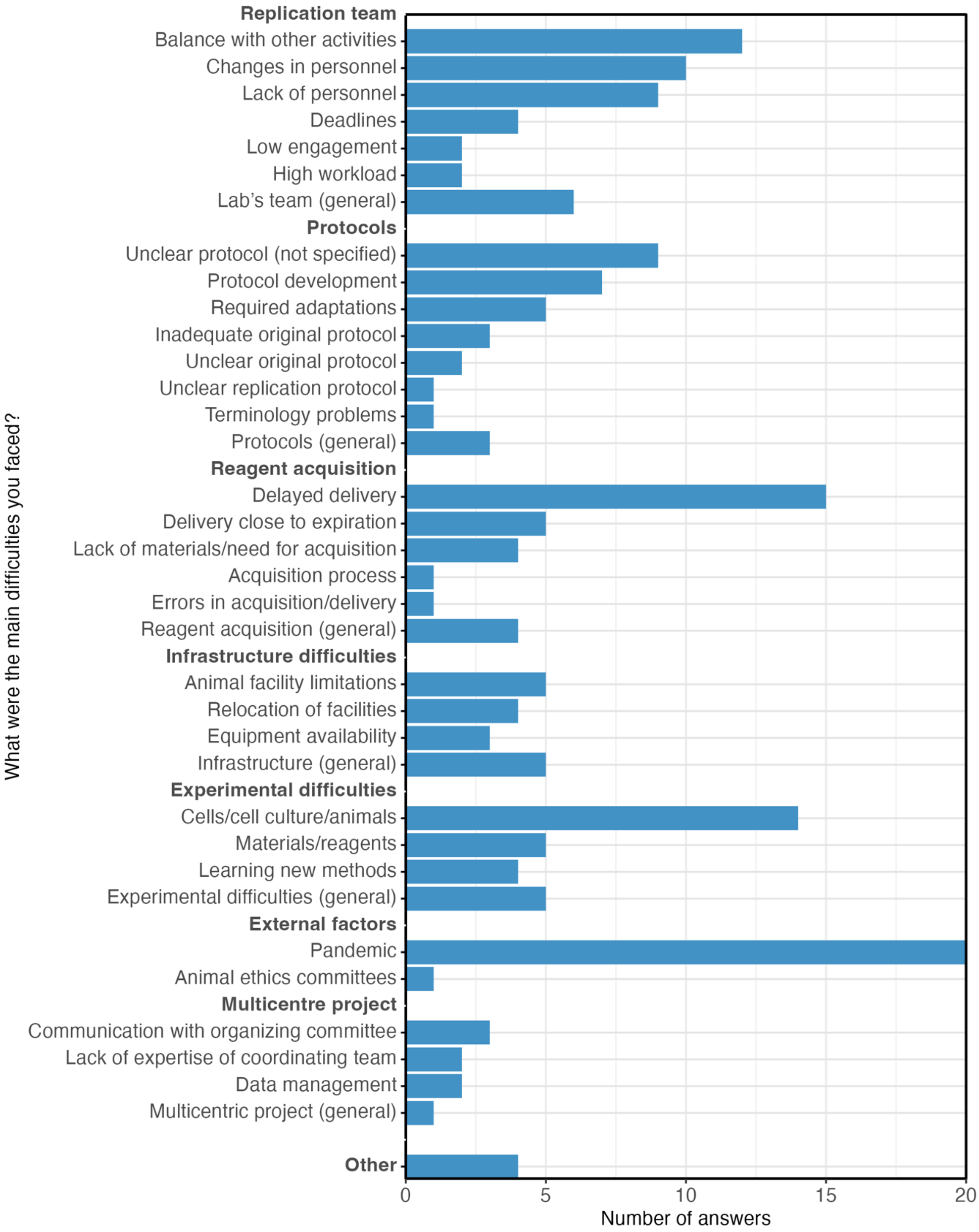


**Figure S9.** Main difficulties faced by labs throughout the project. Reasons were provided as open answers to an anonymous survey answered by 121 project participants (see https://osf.io/nfr6y) and categorized by the coordinating team as described in https://osf.io/5gjb7. Examples for each category are shown in **Table S22**.

| | Registered<br>N = 97 | Included<br>N = 75 | Contributed Data<br>N = 56 |
|---|---|---|---|
| **Region** | | | |
| Southeast | 60 (62%) | 46 (61%) | 36 (64%) |
| South | 19 (20%) | 17 (23%) | 16 (29%) |
| Northeast | 11 (11%) | 9 (12%) | 3 (5.4%) |
| North | 4 (4.1%) | 2 (2.7%) | 0 (0%) |
| Central-West | 3 (3.1%) | 1 (1.3%) | 1 (1.8%) |
| **State** | | | |
| Rio de Janeiro | 28 (29%) | 20 (27%) | 16 (29%) |
| São Paulo | 22 (23%) | 16 (21%) | 12 (21%) |
| Rio Grande do Sul | 9 (9.3%) | 9 (12%) | 9 (16%) |
| Minas Gerais | 9 (9.3%) | 9 (12%) | 7 (13%) |
| Santa Catarina | 6 (6.2%) | 4 (5.3%) | 3 (5.4%) |
| Paraná | 4 (4.1%) | 4 (5.3%) | 4 (7.1%) |
| Rio Grande do Norte | 3 (3.1%) | 3 (4.0%) | 0 (0%) |
| Alagoas | 1 (1.0%) | 1 (1.3%) | 1 (1.8%) |
| Bahia | 1 (1.0%) | 1 (1.3%) | 1 (1.8%) |
| Distrito Federal | 1 (1.0%) | 1 (1.3%) | 1 (1.8%) |
| Espírito Santo | 1 (1.0%) | 1 (1.3%) | 1 (1.8%) |
| Paraíba | 1 (1.0%) | 1 (1.3%) | 1 (1.8%) |
| Pernambuco | 2 (2.1%) | 1 (1.3%) | 0 (0%) |
| Amapá | 1 (1.0%) | 1 (1.3%) | 0 (0%) |
| Amazonas | 1 (1.0%) | 1 (1.3%) | 0 (0%) |
| Ceará | 1 (1.0%) | 1 (1.3%) | 0 (0%) |
| Sergipe | 1 (1.0%) | 1 (1.3%) | 0 (0%) |
| Acre | 1 (1.0%) | 0 (0%) | 0 (0%) |
| Goiás | 1 (1.0%) | 0 (0%) | 0 (0%) |
| Maranhão | 1 (1.0%) | 0 (0%) | 0 (0%) |
| Mato Grosso | 1 (1.0%) | 0 (0%) | 0 (0%) |
| Rondônia | 1 (1.0%) | 0 (0%) | 0 (0%) |

**Table S1. Geographic Distribution of Laboratories.** Number (% of total) of labs in each region/state of Brazil for registered laboratories (those registering to participate in the project), included ones (those that received experiments to replicate) and those that concluded at least one experiment and contributed data to the project.

| | All (n=202) | Protocol (n=41) | Data Collection (n=37) | Both (n=124) |
|---|---|---|---|---|
| **Highest degree** | | | | |
| PhD | 128 (63.4%) | 34 (82.9%) | 7 (18.9%) | 87 (70.2%) |
| MSc | 42 (20.8%) | 4 (9.8%) | 15 (40.5%) | 23 (18.5%) |
| Undergraduate | 23 (11.4%) | 3 (7.3%) | 8 (21.6%) | 12 (9.7%) |
| Secondary School | 9 (4.5%) | 0 (0.0%) | 7 (18.9%) | 2 (1.6%) |
| **Year of degree** | 2016 (1974 - 2024) | 2008 (1974 - 2022) | 2021 (2008 - 2024) | 2016 (1983 - 2023) |
| **Years since first article** | 11 (0 - 45) | 21 (0 - 45) | 2 (0 - 16) | 13 (0 - 44) |
| **h-index** | 6 (0 - 43) | 15 (0 - 39) | 0 (0 - 7) | 7 (0 - 43) |
| **Number of articles** | 11 (0 - 251) | 35 (0 - 251) | 1 (0 - 14) | 12 (0 - 243) |
| **Researchers without publications** | 23 (11.4%) | 1 (2.4%) | 18 (48.6%) | 4 (3.2%) |
| **Number of citations** | 116 (0 - 6328) | 657 (0 - 5617) | 0 (0 - 146) | 162 (0 - 6328) |
| **University ranking** | 5 (1 - 154) | 4 (1 - 86) | 5 (1 - 93) | 7 (1 - 154) |

**Table S2. Profile of researchers involved in protocol design and data collection.** Table shows the number (%) of researchers in each category for categorical variables and median (range) for continuous variables. Information is provided for all researchers, as well as for specific authorship categories (protocol design, data collection or both, with most researchers involved in both) as informed by the participating lab. Information on degrees was obtained from Brazil's Lattes database, while bibliometric information was obtained from the Web of Science between May and December 2024. The Folha University Ranking is a rank of Brazilian universities published yearly (https://ruf.folha.uol.com.br/) and rankings refer to the 2023 edition.

| Experiment (n=60) | |
|---|---|
| Biological model | |
| *Cell line* | 35 (58%) |
| *Rats* | 16 (27%) |
| *Mice* | 6 (10%) |
| *Primary culture (mouse cells)* | 3 (5%) |
| Sample size (per group) | 4.3 (2 - 24) |
| Effect size (Hedges's g) | 2.3 (0.9 - 121.3) |
| Coefficient of variation (original) | 0.2 (0.01 - 1) |
| Reported p value | |
| *< 0.05* | 45 (75%) |
| *< 0.01* | 11 (18%) |
| *< 0.001* | 3 (5%) |
| *< 0.0001* | 1 (2%) |
| Calculated p value | 0.008 ($9x10^{-20}$- 0.11) |
| **Bias control (EPM experiments, n=20)** | |
| Randomization | 8 (40%) |
| Blinded/automated outcome assessment | 7 (35%) |
| Sample size calculation | 1 (5%) |
| Inclusion/exclusion criteria | 5 (25%) |
| **Article (n=60)** | |
| Year of publication | 2013 (2001 - 2017) |
| Number of citations (2 years) | 4 (0 - 17) |
| Area-normalized journal impact factor | 1.07 (0.2 - 4.1) |
| **Corresponding author (n=60)** | |
| Years since first article | 19 (4 - 72) |
| Number of articles (5 previous years) | 33 (0 - 315) |
| **Institution (n=60)** | |
| University ranking | 6 (1 - 110) |
| Region | |
| *Southeast* | 31 (52%) |
| *South* | 21 (35%) |
| *Northeast* | 6 (10%) |
| *Central-West* | 2 (3%) |
| Type | |
| *Public university* | 53 (88%) |
| *Private university* | 6 (10%) |
| *Research institute* | 1 (1%) |

**Table S3. Features of selected experiments, articles, authors and institutions.** Features of experiments selected for replication, as well as their respective articles, corresponding authors and institutions (n=60 for all categories). Sample sizes are estimated as the average between upper and lower bounds of ranges when necessary. Effect sizes and coefficients of variation are calculated based on the difference and error bars extracted from graph, estimating standard deviations based on standard error of mean and sample size when necessary. For experiments reporting standard error and a range for sample size, standard deviations were estimated using the mean of the reported range. Reported p values are displayed as in the article, while calculated p values refer to those of a Welch's t test (for independent samples) or one-sample t test (for presumably paired samples – see https://osf.io/wxzr7) using the extracted difference and standard error. Bias control measures refer to animal experiments only, and indicate the prevalence of randomization, blinding, sample size calculation or definition of inclusion/exclusion criteria for any experiment in the selected articles. Experiments with automated assessment of EPM outcomes were counted as blinded. Bibliometric information was obtained from the Web of Science, with article citations referring to the year of online publication and the 2 subsequent years, and numbers of articles by authors referring to the year of online publication and the 5 previous years. The Folha University Ranking is a rank of Brazilian universities published yearly (https://ruf.folha.uol.com.br/) and rankings refer to the 2019 edition. Further information on data collection is available at https://osf.io/enjxy.

| **All replications** (n= 143) | | | | | |
|---|---|---|---|---|---|
| **# of replications** | **3** | **2** | **1** | **0** | **Total** |
| **EPM** | 10 | 4 | 3 | 3 | 20 |
| **MTT** | 10 | 10 | 0 | 0 | 20 |
| **PCR** | 14 | 5 | 0 | 1 | 20 |
| **Total** | 34 | 19 | 3 | 4 | 60 |
| **Validated replications** (n=90) | | | | | |
| **# of replications** | **3** | **2** | **1** | **0** | **Total** |
| **EPM** | 4 | 7 | 3 | 6 | 20 |
| **MTT** | 3 | 10 | 4 | 3 | 20 |
| **PCR** | 2 | 10 | 2 | 6 | 20 |
| **Total** | 9 | 27 | 9 | 15 | 60 |

**Table S4. Number of replications per experiment.** Number of replications per experiment that were performed either completely or partially (top) and that were considered valid independent replications by the validation committee (bottom). An additional five replications (4 MTT, 1 PCR) had insufficient sample size but n > 1 and were included in meta-analyses but not analyzed independently. Cells display the number of experiments for each method with three, two, one or zero independent replications in these two samples.

| Validation decision | # of cases | Lab agreement | Summary description |
| --- | --- | --- | --- |
| Included | 90 | 90 (100%) | Validated experiments |
| Protocol deviation | 18 | 15 (83%) | Deviations from the protocol were judged to be relevant enough to invalidate the experiment as a direct replication. |
| Inadequate experimental unit | 12 | 9 (75%) | Biological variation between experimental units was judged insufficient. These units were considered technical replicates for the primary analysis, which led sample size to be insufficient. |
| Insufficient sample size | 9 | 9 (100%) | Achieved sample size was not enough to provide 80% power to detect the original standardized difference. Experiments could be included in meta-analyses but were not analyzed independently. |
| Lack of documentation | 6 | 2 (33%) | Documentation in the data collection spreadsheet was not adequate to ensure that the protocol was followed correctly. |
| Non-interpretable results | 2 | 1 (50%) | No valid data was obtained due to methodological reasons and/or pre-established validity criteria. |
| Protocol deviation/ non-interpretable results | 3 | 2 (100%) | Experiments invalidated for more than one reason as described above. |
| Protocol deviation/ insufficient sample size | 2 | 2 (100%) | Experiments invalidated for more than one reason as described above. |
| Protocol deviation/ lack of documentation | 1 | 1 (100%) | Experiments invalidated for more than one reason as described above. |
| **Total** | 143 | 131 (92%) | |

**Table S5. Agreement with validation decisions.** Table shows numbers for the different decisions reached at the validation process, as well as the rate of agreement of labs with each of them and a summary description of each category. For more information on the validation process, see https://osf.io/e3fjg.

| Experimental unit | Accepted? |
| --- | --- |
| **Cell lines** | |
| Different passages of the cell culture were used, with experiments performed on different days. | Yes (default) |
| A culture was subcultured into multiple plates that were maintained in parallel for multiple passages until the day of the experiment, when all of them were used. | Yes |
| Multiple cultures were maintained in parallel but were pooled together before the experiment, with the experimental unit consisting of an experimental well directly subcultured from this pool. | No |
| A single culture was used, and the experimental unit consisted of an experimental well directly subcultured from this culture. | No |
| **Primary culture** | |
| Primary cultures obtained from cells of independent animals or pools of animals were used. | Yes (default) |
| Multiple animals were used for obtaining cultures, but cells from all of them were pooled together, with the experimental units consisting of subcultures of this pool. | No |

**Table S6. Validation decisions for different interpretations of the experimental unit.** Protocols drafted by the coordinating team defined the experimental unit as subsequent passages of a cell line, or as primary cultures from independent animals or pools of animals (i.e. the default option presented for each category). Different definitions of the experimental unit used by labs are presented below this option, with the corresponding validation decisions on the right column.

| Category | N (%) | Description and examples |
|---|---|---|
| **Successful Replications (n=44)** | | |
| Similar effect size | 20 (45%) | The replication found an effect of similar or comparable magnitude to the original *"We achieved a reduction in cell viability of 63.86 % (...), compared to 66.8 % in the original study."* |
| Effect in the same direction | 13 (30%) | The replication found an effect of in the same direction of the original, regardless of the magnitude. "*There was greater expression in the capsaicin-treated group on day 10 than in the control in both the article and our experiment*." |
| Statistically significant | 10 (23%) | The replication found a statistically significant difference between groups of interest. *"It is significantly altered in relation to the non-tumor cell, corroborating the finding, although the expression levels found in MDA-MB-435 were not as high as in the article."* |
| Similar results (not specified) | 2 (5%) | The response mentions "similar results" but does not specify how similarity was defined. "*The results obtained by our group were similar to those described in the original paper*." |
| No rationale | 2 (5%) | The response only states the replication was successful, without providing a rationale. |
| **Unsuccessful Replications (n=99)** | | |
| Not statistically significant | 34 (34%) | The replication did not find a statistically significant difference between groups of interest. "*From these data, we could see that there was no significant difference between the control group and the group treated with chlordiazepoxide."* |
| No effect | 20 (20%) | The replication found no difference between groups that would indicate an effect of the intervention tested. "*We calculated the mean and standard error of the mean number of entries in the open arms from the results obtained in the experiment. From this data, it is possible to predict that there was no difference between the groups."* |
| Effect in the opposite direction | 18 (18%) | The replication found an effect in the opposite direction of the original. *"The authors found a reduction of around 20% in the viability of cells (...). In my experiment, (...) I found an increase of 50% (...)."* |

| | | |
|---|---|---|
| Different effect size | 14 (14%) | The replication found an effect in the same direction but of different magnitude compared to the original. *"The relationship between the OD values obtained from the different groups (control and test) was different from that described in the article."* |
| Failure to replicate method | 12 (12%) | The replication result could not be adequately assessed because the method was not successfully replicated. "*The cells did not differentiate into erythrocytes."* |
| High variance | 10 (10%) | The replication results presented high variation, preventing the result from being adequately interpreted or limiting statistical power. *"The PCR of the target gene did not show replicable amplification, showing visible amplification in some samples and not in others."* |
| Low sample size | 4 (4%) | The replication results were based on a low sample size, preventing the result from being adequately interpreted or limiting statistical power. *"The small sample size obtained at the end of the experiments makes any interpretation of the data difficult. (...) the large variability and the small n make our results inconclusive."* |
| Different result (not specified) | 4 (4%) | The response mentions "different results" but does not specify how this difference was defined. "*Comparing our results with the originals, they differ."* |
| No rationale | 6 (6%) | The response only states the replication was not successful, without providing a rationale. |

**Table S7. Rationale for subjective assessment of replication.** Table categorizes replicating labs' responses to justify their subjective assessments of success for individual replications. Descriptions of each category and illustrative examples are provided on the right column. Responses could be included in multiple categories (e.g. similar effect size and statistically significant; not statistically significant due to high variance and low sample size); thus, the sum of all categories exceeds 100%.

| **By experiment** | **All** | **MTT** | **PCR** | **EPM** |
|---|---|---|---|---|
| Original in replication's 95% PI | **13/34 (38%)** | 7/14 (50%) | 5/9 (56%) | 1/11 (9%) |
| Replication in original 95% CI | **11/35 (31%)** | 4/14 (29%) | 4/10 (40%) | 3/11 (27%) |
| Same-sign significance (p<0.05) | **9/35 (26%)** | 4/14 (29%) | 3/10 (30%) | 2/11 (18%) |
| ≥50% replications significant | **9/35 (26%)** | 4/14 (29%) | 3/10 (30%) | 2/11 (18%) |
| ≥50% subjectively replicated | **12/35 (34%)** | 5/14 (36%) | 4/10 (40%) | 3/11 (27%) |
| **By replication** | **All** | **MTT** | **PCR** | **EPM** |
| Replication in original 95% CI | **21/76 (28%)** | 6/29 (21%) | 5/21 (24%) | 10/26 (38%) |
| Same-sign significance (p<0.05) | **16/76 (21%)** | 9/29 (31%) | 5/21 (24%) | 2/26 (8%) |
| Subjectively replicated | **21/76 (28%)** | 9/29 (31%) | 7/21 (33%) | 5/26 (19%) |

**Table S8. Replication rates for experiments with ≥ 80% *a posteriori* power.** Replication rates when only experiments with 80% *a posteriori* power (calculated for the aggregate of replications using the original relative difference and the variability observed in replications) are considered. Power calculations are performed by simulations, using the original effect size and the individual sample size and standard deviation found in each replication (see https://osf.io/9hj7t). Same-sign significance is based on a fixed meta-analysis estimate, while effect size comparisons are based on random-effects meta-analysis. The 95% prediction interval criterion only uses experiments with more than one replication and thus has a different sample size. All statistical tests use t distributions based on the number of experimental units. PI, prediction interval; CI, confidence interval; MTT, (3-[4,5-dimethylthiazol-2-yl]-2,5 diphenyl tetrazolium bromide) assay; PCR, reverse transcription polymerase chain reaction; EPM, elevated plus maze. For more information on replication criteria, see https://osf.io/9rnuj.

| **By experiment** | **All** | **MTT** | **PCR** | **EPM** |
|---|---|---|---|---|
| Original in replication's 95% PI | **25/52 (48%)** | 11/19 (58%) | 12/19 (63%) | 2/14 (14%) |
| Replication in original 95% CI | **14/56 (25%)** | 5/20 (25%) | 5/19 (26%) | 4/17 (24%) |
| Same-sign significance ($p<0.05$) | **12/56 (21%)** | 7/20 (35%) | 4/19 (21%) | 1/17 (6%) |
| ≥50% replications significant | **10/56 (18%)** | 6/20 (30%) | 4/19 (21%) | 0/17 (0%) |
| ≥50% subjectively replicated | **19/56 (34%)** | 9/20 (45%) | 8/19 (42%) | 2/17 (12%) |
| **By replication** | **All** | **MTT** | **PCR** | **EPM** |
| Replication in original 95% CI | **35/136 (26%)** | 9/48 (19%) | 13/48 (27%) | 13/40 (32%) |
| Same-sign significance ($p<0.05$) | **23/136 (17%)** | 14/48 (29%) | 7/48 (15%) | 2/40 (5%) |
| Subjectively replicated | **41/136 (30%)** | 18/48 (38%) | 17/48 (35%) | 6/40 (15%) |

**Table S9. Replication rates considering all experiments.** Replication rates considering all experiments, irrespective of validation decision or sample size. Experimental units for analysis are also defined by the replicating lab, irrespective of the validation committee's definition. Same-sign significance is based on a fixed meta-analysis estimate, while effect size comparisons are based on random-effects meta-analysis. The 95% prediction interval criterion only uses experiments with more than one replication and thus has a different sample size. All statistical tests use t distributions based on the number of experimental units. PI, prediction interval; CI, confidence interval; MTT, (3-[4,5-dimethylthiazol-2-yl]-2,5 diphenyl tetrazolium bromide) assay; PCR, reverse transcription polymerase chain reaction; EPM, elevated plus maze. For more information on replication criteria, see https://osf.io/9rnuj.

| By experiment | All | MTT | PCR | EPM |
|---|---|---|---|---|
| Original in replication's 95% PI | **22/43 (51%)** | 11/17 (65%) | 9/14 (64%) | 2/12 (17%) |
| Replication in original 95% CI | **15/56 (27%)** | 6/20 (30%) | 5/19 (26%) | 4/17 (24%) |
| Same-sign significance (p<0.05) | **13/56 (23%)** | 7/20 (35%) | 5/19 (26%) | 1/17 (6%) |
| ≥50% replications significant | **11/56 (20%)** | 6/20 (30%) | 4/19 (21%) | 1/17 (6%) |
| ≥50% subjectively replicated | **19/56 (34%)** | 8/20 (40%) | 8/19 (42%) | 3/17 (18%) |
| **By replication** | **All** | **MTT** | **PCR** | **EPM** |
| Replication in original 95% CI | **31/116 (27%)** | 9/45 (20%) | 12/37 (32%) | 10/34 (29%) |
| Same-sign significance (p<0.05) | **22/116 (19%)** | 14/45 (31%) | 6/37 (16%) | 2/34 (6%) |
| Subjectively replicated | **35/116 (30%)** | 17/45 (38%) | 13/37 (35%) | 5/34 (15%) |

**Table S10. Replication rates considering all experiments judged valid by labs.** Replication rates considering all experiments initially judged valid by the lab, irrespective of validation decision. These initial assessments may differ from those informed by labs after receiving the validation committee's decision. Experimental units for analysis are also defined by the replicating lab, irrespective of the validation committee's definition. Same-sign significance is based on a fixed meta-analysis estimate, while effect size comparisons are based on random-effects meta-analysis. The 95% prediction interval criterion only uses experiments with more than one replication and thus has a different sample size. All statistical tests use t distributions based on the number of experimental units. PI, prediction interval; CI, confidence interval; MTT, (3-[4,5-dimethylthiazol-2-yl]-2,5 diphenyl tetrazolium bromide) assay; PCR, reverse transcription polymerase chain reaction; EPM, elevated plus maze. For more information on replication criteria, see https://osf.io/9rnuj.

| By experiment | All | MTT | PCR | EPM |
|---|---|---|---|---|
| Original in replication's 95% PI | **16/36 (44%)** | 8/13 (62%) | 7/12 (58%) | 1/11 (9%) |
| Replication in original 95% CI | **11/36 (31%)** | 5/13 (38%) | 3/12 (25%) | 3/11 (27%) |
| Same-sign significance (p<0.05) | **9/36 (25%)** | 4/13 (31%) | 3/12 (25%) | 2/11 (18%) |
| ≥50% replications significant | **10/36 (28%)** | 5/13 (38%) | 3/12 (25%) | 2/11 (18%) |
| ≥50% subjectively replicated | **15/36 (42%)** | 6/13 (46%) | 6/12 (50%) | 3/11 (27%) |
| **By replication** | **All** | **MTT** | **PCR** | **EPM** |
| Replication in original 95% CI | **21/81 (26%)** | 7/29 (24%) | 4/26 (15%) | 10/26 (38%) |
| Same-sign significance (p<0.05) | **17/81 (21%)** | 10/29 (34%) | 5/26 (19%) | 2/26 (8%) |
| Subjectively replicated | **25/81 (31%)** | 10/29 (34%) | 10/26 (38%) | 5/26 (19%) |

**Table S11. Replication rates for experiments with at least 2 valid replications.** Replication rates considering only experiments with at least 2 valid replications. Same-sign significance is based on a fixed meta-analysis estimate, while effect size comparisons are based on random-effects meta-analysis. The 95% prediction interval criterion only uses experiments with more than one replication and thus has a different sample size. All statistical tests use t distributions based on the number of experimental units. PI, prediction interval; CI, confidence interval; MTT, (3-[4,5-dimethylthiazol-2-yl]-2,5 diphenyl tetrazolium bromide) assay; PCR, reverse transcription polymerase chain reaction; EPM, elevated plus maze. For more information on replication criteria, see https://osf.io/9rnuj.

| By experiment | All | MTT | PCR | EPM |
|---|---|---|---|---|
| Original in replication's 95% PI | **4/9 (44%)** | 2/3 (67%) | 2/2 (100%) | 0/4 (0%) |
| Replication in original 95% CI | **4/9 (44%)** | 1/3 (33%) | 1/2 (50%) | 2/4 (50%) |
| Same-sign significance ($p<0.05$) | **2/9 (22%)** | 1/3 (33%) | 1/2 (50%) | 0/4 (0%) |
| ≥50% replications significant | **1/9 (11%)** | 1/3 (33%) | 0/2 (0%) | 0/4 (0%) |
| ≥50% subjectively replicated | **3/9 (33%)** | 1/3 (33%) | 1/2 (50%) | 1/4 (25%) |
| **By replication** | **All** | **MTT** | **PCR** | **EPM** |
| Replication in original 95% CI | **11/27 (41%)** | 2/9 (22%) | 2/6 (33%) | 7/12 (58%) |
| Same-sign significance ($p<0.05$) | **4/27 (15%)** | 3/9 (33%) | 1/6 (17%) | 0/12 (0%) |
| Subjectively replicated | **9/27 (33%)** | 3/9 (33%) | 3/6 (50%) | 3/12 (25%) |

**Table S12. Replication rates for experiments with 3 valid replications.** Replication rates considering only experiments with 3 valid replications. Same-sign significance is based on a fixed meta-analysis estimate, while effect size comparisons are based on random-effects meta-analysis. The 95% prediction interval criterion only uses experiments with more than one replication and thus has a different sample size. All statistical tests use t distributions based on the number of experimental units. PI, prediction interval; CI, confidence interval; MTT, (3-[4,5-dimethylthiazol-2-yl]-2,5 diphenyl tetrazolium bromide) assay; PCR, reverse transcription polymerase chain reaction; EPM, elevated plus maze. For more information on replication criteria, see https://osf.io/9rnuj.

| By experiment | All | MTT | PCR | EPM |
|---|---|---|---|---|
| Original in replication's 95% PI | **16/39 (41%)** | 8/15 (53%) | 7/13 (54%) | 1/11 (9%) |
| Replication in original 95% CI | **9/45 (20%)** | 4/17 (24%) | 2/14 (14%) | 3/14 (21%) |
| Same-sign significance ($p<0.05$) | **14/45 (31%)** | 7/17 (41%) | 5/14 (36%) | 2/14 (14%) |
| ≥50% replications significant | **12/45 (27%)** | 7/17 (41%) | 3/14 (21%) | 2/14 (14%) |
| ≥50% subjectively replicated | **19/45 (42%)** | 8/17 (47%) | 7/14 (50%) | 4/14 (29%) |
| **By replication** | **All** | **MTT** | **PCR** | **EPM** |
| Replication in original 95% CI | **15/90 (17%)** | 4/33 (12%) | 4/28 (14%) | 7/29 (24%) |
| Same-sign significance ($p<0.05$) | **20/90 (22%)** | 13/33 (39%) | 5/28 (18%) | 2/29 (7%) |
| Subjectively replicated | **29/90 (32%)** | 12/33 (36%) | 11/28 (39%) | 6/29 (21%) |

**Table S13. Replication rates in the primary analysis (z distribution).** Replication rates for the primary analysis using multiple criteria. Results are the same as in **Table 1**, but using z distributions for statistical tests, confidence and prediction intervals (for meta-analyses, individual replications and original results). Same-sign significance is based on a fixed meta-analysis estimate, while effect size comparisons are based on random-effects meta-analysis. The 95% prediction interval criterion only uses experiments with more than one replication and thus has a different sample size. PI, prediction interval; CI, confidence interval; MTT, (3-[4,5-dimethylthiazol-2-yl]-2,5 diphenyl tetrazolium bromide) assay; PCR, reverse transcription polymerase chain reaction; EPM, elevated plus maze. For more information on replication criteria, see https://osf.io/9rnuj.

| By experiment | Primary | Lab's choice | All Exps | ≥ 2 copies | 3 copies | ≥80% power |
|---|---|---|---|---|---|---|
| Original in replication's 95% PI | **16/39 (41%)** | 21/43 (49%) | 23/52 (44%) | 15/36 (42%) | 4/9 (44%) | 14/37 (38%) |
| Replication in original 95% CI | **9/45 (20%)** | 9/56 (16%) | 8/56 (14%) | 8/36 (22%) | 4/9 (44%) | 8/39 (21%) |
| Same-sign significance (p<0.05) | **14/45 (31%)** | 16/56 (29%) | 15/56 (27%) | 13/36 (36%) | 3/9 (33%) | 13/39 (33%) |
| ≥50% replications significant | **12/45 (27%)** | 14/56 (25%) | 13/56 (23%) | 11/36 (31%) | 1/9 (11%) | 11/39 (28%) |
| ≥50% subjectively replicated | **19/45 (42%)** | 19/56 (34%) | 19/56 (34%) | 15/36 (42%) | 3/9 (33%) | 16/39 (41%) |
| **By replication** | **Primary** | **Lab's choice** | **All Exps** | **≥ 2 copies** | **3 copies** | **80% power** |
| Replication in original 95% CI | **15/90 (17%)** | 21/116 (18%) | 22/136 (16%) | 14/81 (17%) | 9/27 (33%) | 14/83 (17%) |
| Same-sign significance (p<0.05) | **20/90 (22%)** | 27/116 (23%) | 28/136 (21%) | 19/81 (23%) | 5/27 (19%) | 19/83 (23%) |
| Subjectively replicated | **29/90 (32%)** | 35/116 (30%) | 41/136 (30%) | 25/81 (31%) | 9/27 (33%) | 26/83 (31%) |

**Table S14. Replication rates for different analysis sets (z distribution).** Replication rates for the primary and secondary analyses. Results are the same as in **Table 2**, but using z distributions for statistical tests, confidence and prediction intervals (for meta-analyses, individual replications and original results). Same-sign significance is based on a fixed meta-analysis estimate, while effect size comparisons are based on random-effects meta-analysis. Subsets for secondary analyses include all experiments judged valid by the replicating lab (Lab's choice), all concluded experiments (All Exps) – both of which use the experimental unit as defined by the lab rather than by the validation committee –, only experiments with at least 2 (≥ 2 Reps) and 3 replications (3 Reps), and only experiments with ≥ 80% *a posteriori* power using the original relative difference and the variability achieved in replications (considering a z distribution, which leads to a different subset than that included in **Table 2** and **Table S8**). The 95% prediction interval criterion only uses experiments with more than one replication and thus has a different sample size in some analyses. PI, prediction interval; CI, confidence interval; For more information on replication criteria, see https://osf.io/9rnuj.

| By experiment | All | MTT | PCR | EPM |
|---|---|---|---|---|
| Original in replication's 95% PI | **26/39 (67%)** | 12/15 (80%) | 8/13 (62%) | 6/11 (55%) |
| Replication in original 95% CI | **13/45 (29%)** | 5/17 (29%) | 5/14 (36%) | 3/14 (21%) |
| Same-sign significance (p<0.05) | **0/45 (0%)** | 0/17 (0%) | 0/14 (0%) | 0/14 (0%) |
| ≥50% replications significant | **10/45 (22%)** | 5/17 (29%) | 3/14 (21%) | 2/14 (14%) |
| ≥50% subjectively replicated | **19/45 (42%)** | 8/17 (47%) | 7/14 (50%) | 4/14 (29%) |
| **By replication** | **All** | **MTT** | **PCR** | **EPM** |
| Replication in original 95% CI | **23/90 (26%)** | 7/33 (21%) | 6/28 (21%) | 10/29 (34%) |
| Same-sign significance (p<0.05) | **17/90 (19%)** | 10/33 (30%) | 5/28 (18%) | 2/29 (7%) |
| Subjectively replicated | **29/90 (32%)** | 12/33 (36%) | 11/28 (39%) | 6/29 (21%) |

**Table S15. Replication rates in the primary analysis (Knapp-Hartung method).** Replication rates for the primary analysis using the Knapp-Hartung method, which uses the number of studies to determine the degrees of freedom for statistical tests. Results are the same as in **Table 1**, but using a t distribution based on the Knapp-Hartung method for statistical tests and prediction intervals derived from meta-analyses. Same-sign significance is based on a fixed meta-analysis estimate, while effect size comparisons are based on random-effects meta-analysis. The 95% prediction interval criterion only uses experiments with more than one replication and thus has a different sample size. PI, prediction interval; CI, confidence interval; MTT, (3-[4,5-dimethylthiazol-2-yl]-2,5 diphenyl tetrazolium bromide) assay; PCR, reverse transcription polymerase chain reaction; EPM, elevated plus maze. For more information on replication criteria, see https://osf.io/9rnuj.

| By experiment | Primary | Lab's choice | All Exps | ≥ 2 copies | 3 copies | ≥80% power |
|---|---|---|---|---|---|---|
| Original in replication's 95% PI | **26/39 (67%)** | 31/43 (72%) | 36/52 (69%) | 25/36 (69%) | 7/9 (78%) | 9/12 (75%) |
| Replication in original 95% CI | **13/45 (29%)** | 15/56 (27%) | 14/56 (25%) | 11/36 (31%) | 4/9 (44%) | 4/13 (31%) |
| Same-sign significance (p<0.05) | **0/45 (0%)** | 3/56 (5%) | 1/56 (2%) | 0/36 (0%) | 0/9 (0%) | 0/13 (0%) |
| ≥50% replications significant | **10/45 (22%)** | 11/56 (20%) | 10/56 (18%) | 10/36 (28%) | 1/9 (11%) | 5/13 (38%) |
| ≥50% subjectively replicated | **19/45 (42%)** | 19/56 (34%) | 19/56 (34%) | 15/36 (42%) | 3/9 (33%) | 6/13 (46%) |
| **By replication** | **Primary** | **Lab's choice** | **All Exps** | **≥ 2 copies** | **3 copies** | **≥80% power** |
| Replication in original 95% CI | **23/90 (26%)** | 31/116 (27%) | 35/136 (26%) | 21/81 (26%) | 11/27 (41%) | 6/30 (20%) |
| Same-sign significance (p<0.05) | **17/90 (19%)** | 22/116 (19%) | 23/136 (17%) | 17/81 (21%) | 4/27 (15%) | 10/30 (33%) |
| Subjectively replicated | **29/90 (32%)** | 35/116 (30%) | 41/136 (30%) | 25/81 (31%) | 9/27 (33%) | 10/30 (33%) |

**Table S16 - Replication rates for different analysis sets (Knapp-Hartung method).** Replication rates for the primary and secondary analyses using the Knapp-Hartung method, which uses the number of studies to determine the degrees of freedom for statistical tests. Results are the same as in **Table 2**, but using a t distribution based on the Knapp-Hartung method for statistical tests and prediction intervals derived from meta-analysis. Same-sign significance is based on a fixed meta-analysis estimate, while effect size comparisons are based on random-effects meta-analysis. Subsets for secondary analyses include all experiments judged valid by the replicating lab (Lab's Choice), all concluded experiments – both of which use the experimental unit as defined by the lab rather than by the validation committee –, only experiments with at least 2 (≥ 2 Reps) and 3 replications (3 Reps), and only experiments with ≥ 80% *a posteriori* power using the original relative difference and the variability achieved in replications (considering the Knapp-Hartung approach, which leads to a different subset than that included in **Table 2** and **Table S8**). The 95% prediction interval criterion only uses experiments with more than one replication and thus has a different sample size. PI, prediction interval, CI, confidence interval. For more information on replication criteria, see https://osf.io/9rnuj.

| By experiment | All | MTT | PCR | EPM |
|---|---|---|---|---|
| Original in replication's 95% PI | **NA** | NA | NA | NA |
| Replication in original 95% CI | **11/45 (24%)** | 5/17 (29%) | 3/14 (21%) | 3/14 (21%) |
| Same-sign significance (p<0.05) | **8/45 (18%)** | 5/17 (29%) | 2/14 (14%) | 1/14 (7%) |
| ≥50% replications significant | **10/45 (22%)** | 5/17 (29%) | 3/14 (21%) | 2/14 (14%) |
| ≥50% subjectively replicated | **19/45 (42%)** | 8/17 (47%) | 7/14 (50%) | 4/14 (29%) |
| **By replication** | **All** | **MTT** | **PCR** | **EPM** |
| Replication in original 95% CI | **23/90 (26%)** | 7/33 (21%) | 6/28 (21%) | 10/29 (34%) |
| Same-sign significance (p<0.05) | **17/90 (19%)** | 10/33 (30%) | 5/28 (18%) | 2/29 (7%) |
| Subjectively replicated | **29/90 (32%)** | 12/33 (36%) | 11/28 (39%) | 6/29 (21%) |

**Table S17. Replication rates in the primary analysis (single mean).** Replication rates for the primary analysis using a single mean. Results are the same as in **Table 1**, but using the mean of all experimental units (normalized by their respective control unit in paired experiments or by the control mean of the respective replication in unpaired ones) rather than a meta-analysis to aggregate results. The prediction interval criterion does not apply for this analysis, as it requires a meta-analysis of multiple replications. PI, prediction interval; CI, confidence interval; MTT, (3-[4,5-dimethylthiazol-2-yl]-2,5 diphenyl tetrazolium bromide) assay; PCR, reverse transcription polymerase chain reaction; EPM, elevated plus maze. For more information on replication criteria, see https://osf.io/9rnuj.

| **By experiment** | **Primary** | **Lab's choice** | **All Exps** | **≥ 2 copies** | **3 copies** | **≥80% power** |
|---|---|---|---|---|---|---|
| Original in replication's 95% PI | **NA** | NA | NA | NA | NA | NA |
| Replication in original 95% CI | **11/45 (24%)** | 15/56 (27%) | 16/56 (29%) | 10/36 (28%) | 4/9 (44%) | 9/28 (32%) |
| Same-sign significance ($p<0.05$) | **8/45 (18%)** | 12/56 (21%) | 12/56 (21%) | 8/36 (22%) | 3/9 (33%) | 7/28 (25%) |
| ≥50% replications significant | **10/45 (22%)** | 11/56 (20%) | 10/56 (18%) | 10/36 (28%) | 1/9 (11%) | 6/28 (21%) |
| ≥50% subjectively replicated | **19/45 (42%)** | 19/56 (34%) | 19/56 (34%) | 15/36 (42%) | 3/9 (33%) | 11/28 (39%) |
| **By replication** | **Primary** | **Lab's choice** | **All Exps** | **≥ 2 copies** | **3 copies** | **≥80% power** |
| Replication in original 95% CI | **23/90 (26%)** | 31/116 (27%) | 35/136 (26%) | 21/81 (26%) | 11/27 (41%) | 19/61 (31%) |
| Same-sign significance ($p<0.05$) | **17/90 (19%)** | 22/116 (19%) | 23/136 (17%) | 17/81 (21%) | 4/27 (15%) | 12/61 (20%) |
| Subjectively replicated | **29/90 (32%)** | 35/116 (30%) | 41/136 (30%) | 25/81 (31%) | 9/27 (33%) | 19/61 (31%) |

**Table S18. Replication rates for different analysis sets (single mean).** Replication rates for the primary and secondary analyses using a single mean. Results are the same as in **Table 2**, but using the mean of all experimental units (normalized by experiment) rather than a meta-analysis to aggregate results. The prediction interval criterion does not apply for this analysis, as it requires a meta-analysis of multiple replications. Subsets for secondary analyses include all experiments judged valid by the replicating lab (Lab's Choice), all concluded experiments – both of which use the experimental unit as defined by the lab rather than by the validation committee –, only experiments with at least 2 (≥ 2 Reps) and 3 replications (3 Reps), and only experiments with ≥ 80% *a posteriori* power using the original relative difference and the variability achieved in replications (considering the single mean approach, which leads to a different subset than that included in **Table 2** and **Table S8**). PI, prediction interval, CI, confidence interval. For more information on replication criteria, see https://osf.io/9rnuj.

| | Primary analysis | | All experiments | |
|---|---|---|---|---|
| **By experiment** | **Paired** | **Original** | **Paired** | **Original** |
| Original in replication's 95% PI | **8/15 (53%)** | 9/15 (60%) | 11/19 (58%) | 12/19 (63%) |
| Replication in original 95% CI | **5/17 (29%)** | 5/17 (29%) | 5/20 (25%) | 5/20 (25%) |
| Same-sign significance ($p<0.05$) | **4/17 (24%)** | 3/17 (18%) | 7/20 (35%) | 6/20 (30%) |
| ≥50% replications significant | **5/17 (29%)** | 5/17 (29%) | 6/20 (30%) | 5/20 (25%) |
| ≥50% subjectively replicated | **8/17 (47%)** | 8/17 (47%) | 9/20 (45%) | 9/20 (45%) |

| | Primary analysis | | All experiments | |
|---|---|---|---|---|
| **By replication** | **Paired** | **Original** | **Paired** | **Original** |
| Replication in original 95% CI | **7/33 (21%)** | 8/33 (24%) | 9/48 (19%) | 9/48 (19%) |
| Same-sign significance ($p<0.05$) | **10/33 (30%)** | 8/33 (24%) | 14/48 (29%) | 12/48 (25%) |
| Subjectively replicated | **12/33 (36%)** | 12/33 (36%) | 18/48 (38%) | 18/48 (38%) |
| Coefficient of variation (median) | **0.11** (0.03-0.92) | 0.30 (0.04-1.47) | 0.11 (0.004-0.92) | 0.25 (0.004-1.47) |

**Table S19. Replication rates for MTT experiments using different approaches for pairing.** "Paired" columns show results of analysis performing paired normalization in every experiment, while "Original" columns show results using the originally registered approach (i.e. pairing only was this was presumably done in the original article). Effect size comparisons are based on random-effects meta-analyses, while same-sign significance is based on fixed-effects meta-analysis estimates. The 95% prediction interval criterion only uses experiments with more than one replication and thus has a different sample size. All statistical tests use the t distributions based on the number of experimental units. Coefficient of variation rows show median and ranges obtained in each analysis. PI, prediction interval, CI, confidence interval. For more information on replication criteria, see the registered replication protocol at https://osf.io/9rnuj.

| | Primary analysis | | All experiments | |
|---|---|---|---|---|
| **By experiment** | **Log values** | **Linear values** | **Log values** | **Linear values** |
| Original in replication's 95% PI | **8/13 (62%)** | 10/13 (77%) | 12/19 (63%) | 16/19 (84%) |
| Replication in original 95% CI | **5/14 (36%)** | 2/14 (14%) | 5/19 (26%) | 5/19 (26%) |
| Same-sign significance (p<0.05) | **3/14 (21%)** | 2/14 (14%) | 4/19 (21%) | 3/19 (16%) |
| ≥50% replications significant | **3/14 (21%)** | 3/14 (21%) | 4/19 (21%) | 4/19 (21%) |
| ≥50% subjectively replicated | **7/14 (50%)** | 7/14 (50%) | 8/19 (42%) | 8/19 (42%) |
| | **Primary analysis** | | **All experiments** | |
| **By replication** | **Log values** | **Linear values** | **Log values** | **Linear values** |
| Replication in original 95% CI | **6/28 (21%)** | 6/28 (21%) | 13/48 (27%) | 12/48 (25%) |
| Same-sign significance (p<0.05) | **5/28 (18%)** | 5/28 (18%) | 7/48 (15%) | 7/48 (15%) |
| Subjectively replicated | **11/28 (39%)** | 11/28 (39%) | 17/48 (35%) | 17/48 (35%) |

**Table S20. Replication rates for PCR experiments using different aggregation methods.** "Log values" columns show results of analysis aggregating ΔCt values for real-time PCR or relative expression in $log_2$ scale for conventional PCR, as in the primary analysis. "Linear values" columns show results aggregating linearized values (usually $2^{-\Delta\Delta Ct}$) for real-time PCR and relative expression in linear scale for conventional PCR, as originally planned in the protocols. Effect size comparisons are based on random-effects meta-analyses, while same-sign significance is based on fixed-effects meta-analysis estimates. The 95% prediction interval criterion only uses experiments with more than one replication and thus has a different sample size. All statistical tests use the t distributions based on the number of experimental units. PI, prediction interval, CI, confidence interval. For more information on replication criteria, see the registered replication protocol at https://osf.io/9rnuj.

| | All (n=70) | MTT (n=22) | PCR (n=18) | EPM (n=30) |
|---|---|---|---|---|
| **Age** | 33 (22 - 65) | 34 (23 - 54) | 30 (23 - 49) | 32 (22 - 65) |
| **Gender** | | | | |
| Female | 32 (46%) | 12 (55%) | 11 (61%) | 9 (30%) |
| Male | 37 (53%) | 10 (45%) | 7 (39%) | 20 (67%) |
| Non-binary | 1 (1%) | 0 (0%) | 0 (0%) | 1 (3%) |
| **BRI member** | | | | |
| Yes | 20 (29%) | 10 (45%) | 2 (11%) | 8 (27%) |
| No | 50 (71%) | 12 (55%) | 16 (89%) | 22 (73%) |
| **Current affiliation** | | | | |
| Brazil | 50 (71%) | 19 (86%) | 11 (61%) | 20 (67%) |
| Europe | 16 (23%) | 2 (9%) | 5 (28%) | 9 (30%) |
| North America | 2 (3%) | 1 (5%) | 1 (6%) | 0 (0%) |
| Asia/Oceania | 1 (1%) | 0 (0%) | 1 (6%) | 0 (0%) |
| Not specified | 1 (1%) | 0 (0%) | 0 (0%) | 1 (3%) |
| **Years in research** | 10 (3 - 46) | 14 (4 - 32) | 7 (3 - 30) | 10 (3 - 46) |
| **Highest academic title** | | | | |
| PhD | 48 (69%) | 16 (73%) | 9 (50%) | 23 (77%) |
| Master's | 17 (24%) | 5 (23%) | 7 (39%) | 5 (17%) |
| Bachelor | 5 (7%) | 1 (5%) | 2 (11%) | 2 (7%) |
| **Institutional position** | | | | |
| Professor | 23 (33%) | 9 (41%) | 4 (22%) | 10 (33%) |
| Post-doc/Research associate | 22 (31%) | 6 (27%) | 4 (22%) | 12 (40%) |
| PhD candidate | 17 (24%) | 5 (23%) | 7 (39%) | 5 (17%) |
| Master's candidate | 4 (6%) | 1 (5%) | 1 (6%) | 2 (7%) |
| Technician | 1 (1%) | 0 (0%) | 1 (6%) | 0 (0%) |
| Other | 3 (4%) | 1 (5%) | 1 (6%) | 1 (3%) |
| **Theoretical knowledge of method** | | | | |
| None | 2 (3%) | 1 (5%) | 0 (0%) | 1 (3%) |
| Slight | 2 (3%) | 0 (0%) | 1 (6%) | 1 (3%) |
| Moderate | 16 (23%) | 3 (14%) | 4 (22%) | 9 (30%) |
| Good | 37 (53%) | 13 (59%) | 11 (61%) | 13 (43%) |
| Excellent | 13 (19%) | 5 (23%) | 2 (11%) | 6 (20%) |
| **Practical experience with method** | | | | |
| None | 12 (17%) | 2 (9%) | 3 (17%) | 7 (23%) |
| A little | 10 (14%) | 1 (5%) | 4 (22%) | 5 (17%) |
| A moderate amount | 22 (31%) | 9 (41%) | 2 (11%) | 11 (37%) |
| A lot | 12 (17%) | 3 (14%) | 4 (22%) | 5 (17%) |
| Performs routinely | 14 (20%) | 7 (32%) | 5 (28%) | 2 (7%) |

| | All (n=70) | MTT (n=22) | PCR (n=18) | EPM (n=30) |
|---|---|---|---|---|
| **Statistics knowledge** | | | | |
| None | 0 (0%) | 0 (0%) | 0 (0%) | 0 (0%) |
| Slight | 1 (1%) | 1 (5%) | 0 (0%) | 0 (0%) |
| Moderate | 20 (29%) | 6 (27%) | 9 (50%) | 5 (17%) |
| Good | 37 (53%) | 14 (64%) | 6 (33%) | 17 (57%) |
| Excellent | 12 (17%) | 1 (5%) | 3 (17%) | 8 (27%) |
| **Expected replication rate** | 40 (10 - 95) | 50 (20 - 90) | 40 (20 - 95) | 40 (10 - 95) |
| **Research area** | | | | |
| Life and health sciences | 68 (100%) | 21 (100%) | 18 (100%) | 29 (100%) |

**Table S21. Survey participant information.** Demographic and self-rated knowledge information on survey participants, as collected by the survey. Table shows the number (%) of researchers in each category for categorical variables and median (range) for continuous variables. Information is provided for all researchers, as well as for those predicting results with each individual method. Theoretical knowledge and practical experience both refer to the method in which participants predicted results. Expected replication rate refers to participant's estimate of the general replication rate of published biomedical results. Although the survey was open to participants in any field of research, all participants were from the life and health sciences. A copy of the survey with the specific questions asked can be found at https://osf.io/29mq5.

| Category | Examples |
|---|---|
| **Experimental issues** | |
| Model behaves differently (n = 20) | *"The growth of the cells was not satisfactory, so the protocol was changed according to the ATCC cultivation recommendations."* |
| Experiment works differently (n = 16) | *"The other deviation that we considered important was the problem in dissolving the drug (extended-release capsules) and administering it."* |
| Different cells or animals (n = 12) | *"The animals supplied varied greatly in weight for animals of the same age."* |
| Changes due to pilot (n = 3) | *"In this case, we detected the need to deviate from the original protocol as we began to cultivate the strain in our laboratory. We noticed that to reach the confluence described in the original protocol for treatment, starting from the cell density also described in the protocol, we would need more than the 24 hours originally described."* |
| Unfeasible validity criteria (n = 2) | *"The mean deviation was in (...) compliance with the criteria for inclusion/exclusion of animals that have or have not consumed the drug."* |
| Experimental issues (general) (n = 10) | *"The common deviations in laboratory practices that exist between different laboratories."* |
| **Infrastructure/logistics** | |
| Lab infrastructure (n = 17) | *"We replaced OneStep RT-PCR with Two Steps because we couldn't program the BioRad thermal cycler for the original protocol."* |
| Animal facility limitations (n = 11) | *"The delay in the ethics committee's approval made it difficult to provide animals of the same age as the original protocol, and these animals would have to be requested from the UFRGS breeding center some time in advance."* |
| Regulatory requirements (n = 7) | *"Requirements of CEUA/Unifesp regarding the number of animals/cage, according to the types of cages we have."* |
| Expired reagents (n = 7) | *"Although every care was taken to carry out the protocol, the reagent expired, which may have had an impact on the results."* |
| Deadlines (n = 7) | *"Lack of time to repeat experiments and improve quality."* |
| Pandemic restrictions (n = 6) | *"Most of the deviations occurred because of the pandemic, which limited our access to the laboratory."* |
| Costs (n = 2) | *"We added another control to better evaluate the results and we carried out the MTT test in a 96-well plate, because in a 24-well plate, as indicated, it would have been too costly, with an absurd amount of material"* |
| Infrastructure/logistics (general) (n = 5) | *"Limitations in the working hours of the experimenter."* |

| | |
|---|---|
| **Deliberate choice** | |
| Standard lab protocol<br>(n = 6) | *"Some deviations were caused by the experimenter using the laboratory's "standard protocol" and others by the proximity of the deadline for delivery of the experiment."* |
| Supplier recommendations<br>(n = 6) | *"We didn't use the probe concentration from the protocol because the manufacturer doesn't tell us the exact concentration, but that the reagent is '20x' concentrated, so we used the concentration recommended by the manufacturer."* |
| Team expertise<br>(n = 3) | *"The incubation time of the cells with MTT and the solubilization of the crystals formed with DMSO were adjusted according to our group's previous experience, which clearly indicates that incubation for 1 hour with MTT and solubilization for 10 minutes with DMSO are sufficient to generate absorbance readings within the optimum reading range for the equipment (D.O.>0.1) and pre-saturation."* |
| Deliberate choice (general)<br>(n = 1) | *"The new concentrations of DMSO used were due to the fact that we believed that 2.5% (initial protocol) was too high, which could lead to a change in the final result."* |
| **Lab error** | |
| Error in protocol development<br>(n = 4) | *"The exposure time described in the protocol was not suitable for the number of animals per mini-experiment, which is why the exposure time varied in this way."* |
| Error in planning<br>(n = 4) | *"Mistake in planning the materials to obtain the correct replicates"* |
| Error in executing experiment<br>(n = 3) | *"Inaccurate surgery."* |
| Unclear protocol<br>(n = 3) | *"The methodology was not clearly described at some points, especially in relation to dilution, how the solutions would be given and the lack of control over the exact amount of solution ingested by each animal."* |
| Lab error (general)<br>(n = 3) | *"Incorrect number of cells to be plated."* |
| **Coordinating team error** | |
| Error in acquiring materials<br>(n = 1) | *"The drug was supplied with an excipient when it shouldn't have been."* |
| Error in communication with lab<br>(n = 1) | *"Not knowing in advance how* [the reagent] *would be delivered (solution or powder) also made it difficult to plan the preparation of the solution"* |

**Table S22. Examples of reasons for protocol deviations.** Table shows the various categories and subcategories for protocol deviations (in order of general category frequency, as in **Figure 5**), with illustrative examples of lab responses for each subcategory.

| Category | Examples |
|---|---|
| **Replication team** | |
| Balance with other activities (n = 12) | *"It was also difficult to carry out the experiments for this project at the same time as we had our own projects in progress, to find the best way to manage time throughout the process."* |
| Changes in personnel (n = 10) | *"Participants in the group moved over the course of the projects, which lasted for years."* |
| Lack of personnel (n = 9) | *"(...) conducting complex behavioral assessment experiments with few experimenters involved."* |
| Deadlines (n = 4) | *"As I did everything myself, it took me a while to meet the deadlines."* |
| Low engagement (n = 2) | *"The students' low engagement was disappointing for me. They just did the tasks that were asked of them. They never took part in meetings or other [BRI] activities. Although I explained the importance, I was frustrated by the low proactivity and I think they missed a great learning opportunity."* |
| High workload (n = 2) | *"The commitment throughout the project to respond to demands was more difficult than expected"* |
| Lab's team (general) (n = 6) | *"Difficulty with the other partners in my group"* |
| **Protocols** | |
| Unclear protocol (not specified) (n = 9) | *"(...) to understand some protocols that were not very clear"* |
| Protocol development (n = 7) | *"(...) fill in the gaps in the original protocols with information that made sense from an experimental point of view."* |
| Required adaptations (n = 5) | *"Obtaining the expected results since we went through a change in protocol that directly affected our results."* |
| Inadequate original protocol (n = 3) | *"(...) to follow some experimental designs that might not be suitable for a particular analysis"* |
| Unclear original protocol (n = 2) | *"(...) reproducing a hypoxia experiment that didn't detail the exact concentration of oxygen used in the work."* |
| Unclear replication protocol (n = 1) | *"To understand the protocol, since I took over after this stage."* |
| Terminology problems (n = 1) | *"Misinterpretation of biological and/or technical replicates when developing the protocol."* |
| Protocols (general) (n = 3) | *"Being able to faithfully replicate the original protocol."* |

| | |
|---|---|
| **Reagent acquisition** | |
| Delayed delivery<br>(n = 15) | *"Some difficulties in receiving reagents"* |
| Delivery close to expiration<br>(n = 5) | *"Another issue was the receipt of some expired reagents and culture media, which probably caused changes in the results of some experiments."* |
| Lack of materials/need for acquisition<br>(n = 4) | *"Insufficient materials to correctly carry out the necessary sampling"* |
| Acquisition process<br>(n = 1) | *"Delay in the purchase/arrival of material"* |
| Errors in acquisition/delivery<br>(n = 1) | *"The availability of the material used in an attempt to reproduce the method of a paper. Different reagents (MTT formazan) certainly produce different results."* |
| Reagent acquisition (general)<br>(n = 4) | *"The biggest difficulty was finding the same reagents used in the original article. In my case, the product was discontinued."* |
| **Infrastructure difficulties** | |
| Animal facility limitations<br>(n = 5) | *"Delays and difficulties in (...) the supply of animals (the same age as in the original experiment, etc.)"* |
| Relocation of facilities<br>(n = 4) | *"We also had a change in the culture room which made it take even longer"* |
| Equipment availability<br>(n = 3) | *"In general, we didn't have all the equipment we needed, so we had to go to other academic units at the university."* |
| Infrastructure (general)<br>(n = 5) | *"Lack of electricity in the laboratory which hindered the conduct of the experiments."* |
| **Experimental difficulties** | |
| Cells/cell culture/animals<br>(n = 14) | *"Cell treatment, at first we had a few episodes of contamination."* |
| Materials/reagents<br>(n = 5) | *"Crucial reagents for the endpoints tested without quality control of function."* |
| Learning new methods<br>(n = 4) | *"The main problem encountered was a lack of familiarity with the techniques needed to carry out the experiment (...) culture and differentiation of specific cell lines, treatment with specific factors, stains used to validate some of the previous steps were previously unknown to the group. This made it difficult to deal with the samples and it took some time before the experiment could be reproduced."* |
| Experimental difficulties (general)<br>(n = 5) | *"Thorough adaptation to the proposed protocols"* |

| **External factors** | |
|---|---|
| Pandemic<br>(n = 20) | *“Developing the project during the pandemic.”* |
| Animal ethics committees<br>(n = 1) | *“Initially we faced resistance from our research committee when it came to approving and developing the project.”* |
| **Multicenter project** | |
| Communication with coordinating team<br>(n = 3) | *“I think the main thing is the delay in the initiative's response to our questions and needs.”* |
| Lack of expertise of coordinating team<br>(n = 2) | *“Lack of knowledge of the committee member who worked with us about cell culture (...)”* |
| Data management<br>(n = 2) | *“Organize the data in the spreadsheet so that it is clear, especially the samples that were repeated and excluded.”* |
| Multicenter project (general)<br>(n = 1) | *“Establishing a schedule according to lab member demands and (...) [BRI]'s general progress”* |

**Table S23. Examples of difficulties faced by labs replicating the experiments.** Table shows the various categories and subcategories for general difficulties mentioned in the project evaluation survey (in order of general category frequency, as in **Figure S9**), with illustrative examples of participant responses for each subcategory.

| Category | Difficulty | Rating |
|---|---|---|
| Large-scale project management | Communication difficulties between the coordinating team and participating labs. | 8.2 |
| | Failure to anticipate loss of labs and experiments when planning analyses. | 8.2 |
| | Lack of attention to written guidelines/manuals by participant labs. | 8.2 |
| Researcher practices and expertise | Absence of standardized terminologies to define experimental units and other aspects. | 8.8 |
| | Lack of attention to detail when preparing protocols. | 8.5 |
| | Poor data management. | 8.0 |
| Scientific literature issues | Poorly described methods in original articles. | 8.7 |
| | Poorly described experimental unit in original articles. | 8.7 |
| | Likely underestimation of biological variability in original articles. | 7.8 |
| Intrinsic experimental difficulties | Models behaving differently than expected. | 8.7 |
| | Tension between replicating method as described and replicating model features. | 8.2 |
| | Difficulties in pre-registration without prior testing of the model. | 6.8 |
| Logistics and infrastructure | Issues with materials suppliers, including delays and communication problems. | 8.7 |
| | Difficulties in obtaining licenses with Brazilian regulatory agencies. | 7.8 |
| | Limitations in animal suppliers and facilities. | 7.2 |

**Table S24. Project difficulties - Coordinating team assessment.** Table shows the top 3 difficulties faced by the project in 5 different dimensions, as assessed by independent ratings from 6 members of the coordinating team, with the mean rating – on a scale of 1 (not a relevant difficulty) to 10 (very relevant difficulty with a large impact on the project) – displayed on the right. The full list of difficulties is available at https://osf.io/q76vj.
